# TERRA increases at short telomeres in yeast survivors and regulates survivor associated senescence (SAS)

**DOI:** 10.1101/2022.07.29.502086

**Authors:** Stefano Misino, Anke Busch, Carolin B. Wagner, Brian Luke

## Abstract

Cancer cells achieve immortality by employing either homology-directed repair (HDR) or the telomerase enzyme to maintain telomeres. ALT (alternative lengthening of telomeres) refers to the subset of cancer cells that employ HDR. ALT mechanisms are strongly conserved from yeast to human cells, with the yeast equivalent being referred to as survivors. The non-coding RNA, TERRA and its ability to form RNA-DNA hybrids have been implicated in ALT/survivor maintenance by promoting HDR. It is not understood which telomeres in ALT/survivors engage in HDR, nor is it clear which telomeres upregulate TERRA. Using yeast survivors as a model for ALT, we demonstrate that HDR only occurs at telomeres when they become critically short. Moreover, TERRA levels steadily increase as telomeres shorten and decrease again following HDR-mediated recombination. Surprisingly, we observe that survivors undergo cycles of senescence, in a similar manner to non-survivors following telomerase loss, which we refer to as survivor associated senescence (SAS). Similar to “normal” senescence, we observe that RNA-DNA hybrids slow the rate of SAS by decreasing the rate of telomere shortening. In summary, TERRA RNA-DNA hybrids regulate telomere dysfunction-induced senescence in both pre- and post-crisis cells.

## Introduction

Telomeres are nucleoprotein structures that protect the ends of chromosomes from illegitimate repair and degradation^1^. In budding yeast, *Saccharomyces cerevisiae*, telomeres harbour imperfect double stranded TG1-3 repeats over a length of approximately 350 bp and terminate with a G-rich 3’ overhang of 10-14 nucleotides^2^. The Rap1 protein, along with Rif1, Rif2 and the Sir2/3/4 complex, bind to the dsDNA region, while the CST complex (Cdc13, Stn1 and Ten1) associates with the ssDNA overhang^2,3^. Together, these proteins prevent telomeres from being recognized as DSBs, thereby avoiding repair via non-homologous end-joining (NHEJ) and homology-directed repair (HDR)^1,2,4^, which would ultimately lead to lethal chromosome fusions.

Telomeres shorten with each round of DNA replication due to the end replication problem^5,6,7^. In budding yeast, this occurs at a rate of 3-5 bp per cell division^7^. The progressive telomere attrition leads to the eventual activation of the DNA damage response (DDR) checkpoint and permanent cell cycle arrest. The loss of replicative potential due to telomere shortening is referred to as replicative senescence^3,6,8^. Replicative senescence acts as a tumour suppressor in multi-cellular organisms by preventing uncontrolled cell division; however, the accumulation of senescent cells can contribute to the aging process^9,10^. Hence, rates of telomere shortening must be carefully controlled to ensure a non-pathogenic balance.

In order for cancer cells to overcome replicative senescence and achieve an unlimited replication potential, they must re-establish a telomere maintenance mechanism (TMM). Different strategies have evolved to maintain telomeres and counteract senescence in human cancer cells, the most common being telomerase reactivation. Telomerase is the enzyme specialized in *de novo* synthesis of telomere sequences^3,8,11^. In humans, telomerase is expressed in the early stages of development and then silenced in the majority of somatic tissues^12,13^. 85-90% of human cancers re-express telomerase to bypass replicative senescence and allow unlimited proliferation^14,15,16^. On the other hand, 10-15% of human cancers use an HDR-mediated TMM, that is referred to as the alternative lengthening of telomeres (ALT) mechanism^17,18^.

Similar to humans, budding yeast can employ either telomerase or HDR to maintain its telomeres^8,19,20^. Telomerase is constitutively expressed and active in wild-type cells. When telomerase is deleted, telomeres shorten and cells eventually lose replicative potential as they enter replicative senescence^3,8,19^. Although the majority of cells permanently arrest, a subset of “survivors” employ HDR to elongate their telomeres and escape senescence^20,21^. Two survivor types are known to form: type I and type II survivors. Type I survivors depend on Rad51, Rad54 and Rad57 and present very short telomeres with tandem amplification of the subtelomeric Y’ element. Type II survivors, on the other hand, rely on the Mre11, Rad50, Xrs2 (MRX) complex, Sgs1 and Rad59 and display telomeres with heterogeneous TG1-3 lengths^4,20,21,22^. Due to the common features between type II survivors and ALT human cancers telomeres, this type of survivors has been used to gain mechanistic insights into ALT features. Since type II survivors have a growth advantage over type I survivors, they are selected for when experiments are performed in liquid media, due to competition^21,22^.

Despite the differences between type I and type II survivors, post-senescence maintenance of telomeres is unanimously a break-induced replication (BIR) event that depends on Rad52 and Pol32^20,21,23^. A recent study further supports the idea of a common pathway underlying type I and type II BIR. Accordingly, a unified Rad51-mediated mechanism leads to the initial formation of survivor precursors via recombination, followed by maturation and differentiation into the two survivor types^24^. Some *S. cerevisiae* survivors have been reported to acquire a replicative potential similar to that of wild-type cells and it was presumed that they would eternally maintain a high proliferative potential^20^. However, some survivors did show variations in growth, switching back and forth from fast to slow proliferative rates^20^. This feature seems more evident in type I than in type II survivors^21,22,25^.

TERRA is a long non-coding RNA transcribed at telomeres by RNA polymerase II and degraded by the RNA exonuclease Rat1^26,27^. TERRA forms RNA-DNA hybrids that are removed by the RNase H1 and, primarily, RNase H2 enzymes^28,29^. Both the soluble transcript and the hybrids are regulated in a cell cycle- and telomere length-dependent manner via recruitment of Rat1 and RNase H2 to telomeres, respectively^29^. The function of TERRA is still to be fully elucidated; however, an increasing number of studies show that the transcript is involved in multiple telomeric transactions, including: modulation of telomerase activity, HDR activation, telomeres capping, chromatin regulation and stress signalling^30^. In yeast cells lacking telomerase, TERRA RNA-DNA hybrids accumulate at short telomeres and participate in telomere maintenance^28,29^. Indeed, the accumulation or removal of RNA-DNA hybrids at telomeres can either slow or accelerate rates of replicative senescence, respectively. The precise contribution of TERRA to telomere maintenance remains to be fully elucidated.

In ALT human cancer cells, TERRA is upregulated compared to telomerase-positive cells and dysregulation of TERRA RNA-DNA hybrids compromises telomeres integrity^31–33^. It is thought that TERRA upregulation in these cells derives from less condensed chromatin at telomeres and consequently increased transcription^31,34^. We have previously reported that, similar to human ALT tumors, TERRA accumulates in type II survivors and the abrogation of telomeric R-loops causes growth defects in survivor cells^35^. However, the source of TERRA accumulation in ALT remains elusive, i.e., is TERRA accumulating at all telomeres? Furthermore, it is not clear which telomeres undergo recombination in ALT/survivor cells, i.e., are all telomeres undergoing HDR or only a subset of telomeres?

In this study we demonstrate that, at natural telomeres, type II survivors also undergo telomere attrition over time and experience replicative senescence. We refer to this phenomenon as survivor associated senescence (SAS). Furthermore, we demonstrate that telomeres are not constantly undergoing HDR, but only do so when they become critically short. Strikingly, similar to replicative senescence, SAS is regulated by the presence of TERRA RNA-DNA hybrids. The removal and accumulation of TERRA RNA-DNA hybrids accelerate and slow rates of SAS, respectively. Finally, we observe that in type II survivors TERRA levels are specifically increased when telomeres become critically short, again mirroring what has been reported to occur during replicative senescence.

In conclusion, we report an unexpected parallel between pre- and post-crisis cells with respect to the regulation of telomere maintenance and replicative potential via TERRA. The observation that senescence phenotypes endure beyond immortalization might have important consequences for the treatment of human cancers using ALT as a telomere maintenance mechanism.

## Materials and Methods

### Generation of type II survivors

Diploid cells heterozygous for a telomerase gene (*EST2/est2Δ* or *TLC1*/*tlc1Δ*) were sporulated and dissected. Single clones were isolated and maintained in liquid culture by daily dilution. At the end of senescence, type II survivors were collected and frozen once the replicative potential was regained. For each experiment of this study, survivor cells were streaked to single colonies from cryostocks on appropriate solid media-containing plates and individual clones were further utilized. All yeast strains used in this study can be found in Table 1, and plasmids employed are listed in Table 2.

**Table 1:** Yeast strains used in this study.

| Name | Features | Description |
| --- | --- | --- |
| yDB13 | Wild-type A | *MATa; his3Δ1; leu2Δ0; ura3Δ0; met15Δ0; bar1::KAN + pBL189-URA3* |
| yDB15 | Wild-type B | *MATa; his3Δ1; leu2Δ0; ura3Δ0; met15Δ0; bar1::KAN + pBL189-URA3* |
| yDB05 | Survivor A | *MATa; his3Δ1; leu2Δ0; ura3Δ0; met15Δ0; bar1::KAN; tlc1::HIS3* |
| yDB06 | Survivor B | *MATa; his3Δ1; leu2Δ0; ura3Δ0; met15Δ0; bar1::KAN; tlc1::HIS3* |
| yDB07 | Survivor C | *MATa; his3Δ1; leu2Δ0; ura3Δ0; met15Δ0; bar1::KAN; tlc1::HIS3* |
| yDB08 | Survivor D | *MATa; his3Δ1; leu2Δ0; ura3Δ0; met15Δ0; bar1::KAN; tlc1::HIS3* |
| ySM269 | Wild-type (Rat1 no-TAP) | *MATa; his3Δ1; leu2Δ0; ura3Δ0; met15Δ0* |
| ySM270 | Wild-type (Rat1 no-TAP) | *MATa; his3Δ1; leu2Δ0; ura3Δ0; met15Δ0* |
| yBL7 | Wild-type | *MATa; his3Δ1; leu2Δ0; ura3Δ0; met15Δ0* |
| ySM 272 | Wild-type (Rat1-TAP) | *MATα; his3Δ1; leu2Δ0; ura3Δ0; met15Δ0; rat1-TAP::HIS3* |
| ySM273 | Wild-type (Rat1-TAP) | *MATα; his3Δ1; leu2Δ0; ura3Δ0; met15Δ0; rat1-TAP::HIS3* |
| ySM281 | Survivor (Rat1 no-TAP) | *MATα; his3Δ1; leu2Δ0; ura3Δ0; met15Δ0; tlc1::NAT* |
| ySM282 | Survivor (Rat1 no-TAP) | *MATa; his3Δ1; leu2Δ0; ura3Δ0; met15Δ0; tlc1::NAT* |
| ySM283 | Survivor (Rat1 no-TAP) | *MATa; his3Δ1; leu2Δ0; ura3Δ0; met15Δ0; tlc1::NAT* |
| ySM284 | Survivor (Rat1-TAP) | *MATα; his3Δ1; leu2Δ0; ura3Δ0; met15Δ0; tlc1::NAT; rat1-TAP::HIS3* |
| ySM285 | Survivor (Rat1-TAP) | *MATα; his3Δ1; leu2Δ0; ura3Δ0; met15Δ0; tlc1::NAT; rat1-TAP::HIS3* |
| ySM286 | Survivor (Rat1-TAP) | *MATα; his3Δ1; leu2Δ0; ura3Δ0; met15Δ0; tlc1::NAT; rat1-TAP::HIS3* |
| ySM321 | Survivor (Rnh201 no-TAP) | *his3Δ1; leu2Δ0; ura3Δ0; met15Δ0; est2::NAT* |
| ySM322 | Survivor (Rnh201 no-TAP) | *his3Δ1; leu2Δ0; ura3Δ0; met15Δ0; est2::NAT* |
| ySM326 | Survivor (Rnh201-TAP) | *his3Δ1; leu2Δ0; ura3Δ0; met15Δ0; est2::NAT; rnh201-TAP::HIS* |
| ySM329 | Survivor (Rnh201-TAP) | *his3Δ1; leu2Δ0; ura3Δ0; met15Δ0; est2::NAT; rnh201-TAP::HIS* |
| ySM338 | Wild-type (Rnh201-TAP) | *his3Δ1; leu2Δ0; ura3Δ0; met15Δ0; rnh201-TAP::HIS* |
| ySM342 | Wild-type (Rnh201-TAP) | *his3Δ1; leu2Δ0; ura3Δ0; met15Δ0; rnh201-TAP::HIS* |
| ySM345 | Wild-type (Rnh201 no-TAP) | *his3Δ1; leu2Δ0; ura3Δ0; met15Δ0;* |
| ySM346 | Wild-type (Rnh201 no-TAP) | *his3Δ1; leu2Δ0; ura3Δ0; met15Δ0;* |
| ySM81-86 | Survivor to be transformed with pBL211 (EV) or pBL352 (RNH1) | *MATa; his3Δ1; leu2Δ0; ura3Δ0; met15Δ0; tlc1::HIS* |
| yVP345 | Survivor (*rnh201Δ*) to be transformed with pBL97 (EV) or pBL401 (RNH201) | *MATα; his3Δ1; leu2Δ0; ura3Δ0; met15Δ0; tlc1::NAT; rnh201::HYG* |
| yVP347 | Survivor (*rnh201Δ*) to be transformed with pBL97 (EV) or pBL401 (RNH201) | *MATa; his3Δ1; leu2Δ0; ura3Δ0; met15Δ0; tlc1::NAT; rnh201::HYG* |
| ySM518-530 | Survivor (*rnh201Δ*) to be transformed with pBL97 (EV) or pBL401 (RNH201) | *his3Δ1; leu2Δ0; ura3Δ0; met15Δ0; est2::HYG; rnh201::NAT* |
| ySM123 | yDB05 passaged in liquid culture: 1L telomere length is of ~ 400 bp) | *MATa; his3Δ1; leu2Δ0; ura3Δ0; met15Δ0; bar1::KAN; tlc1::HIS3* |
| yCW450, 454, 458 | Wild-type transformed with pBL906 (EV) | *MATa; his3Δ1; leu2Δ0; ura3Δ0; met15Δ0; osTIR1(F74G)-Leu2 + pBL906 (pRS416)* |
| yCW452, 456, 460 | Wild-type transformed with pBL837 (RNase H1*) | *MATa; his3Δ1; leu2Δ0; ura3Δ0; met15Δ0; osTIR1(F74G)-Leu2 + pBL837 (pRS416 GAL-RNH1-(D193N)-HA)* |
| yCW462, 466, 470 | Survivor transformed with pBL906 (EV) | *MATa; his3Δ1; leu2Δ0; ura3Δ0; met15Δ0; tlc1::NAT; osTIR1(F74G)-Leu2 + pBL906 (pRS416)* |
| yCW464, 468, 472 | Survivor transformed with pBL837 (RNase H1*) | *MATa; his3Δ1; leu2Δ0; ura3Δ0; met15Δ0; tlc1::NAT; osTIR1(F74G)-Leu2 Leu2 + pBL837 (pRS416 GAL-RNH1-(D193N)-HA)* |

**Table 2:**
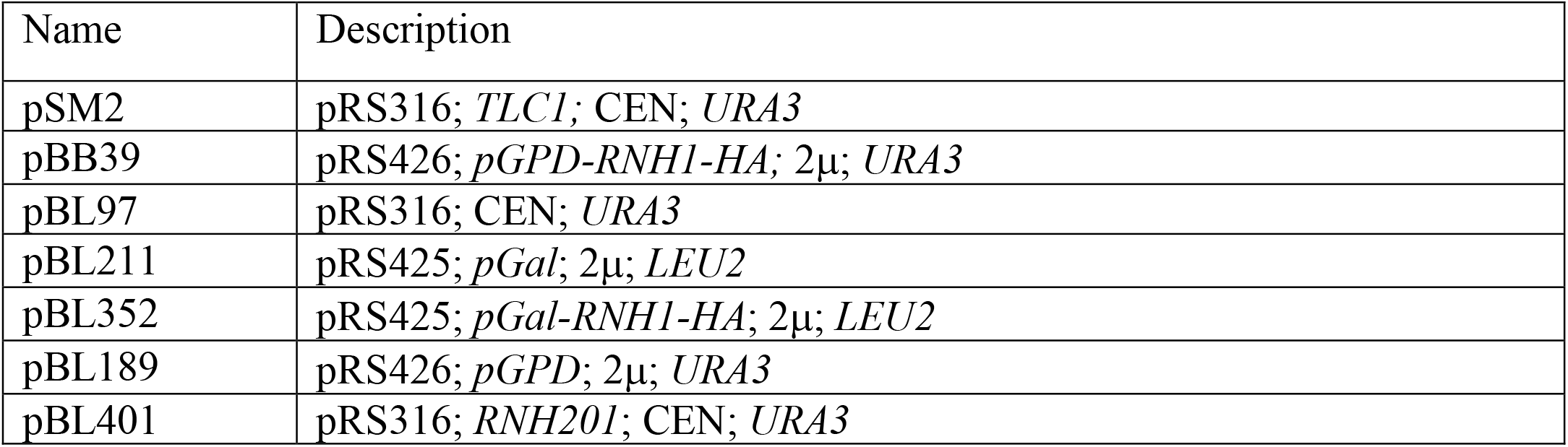
Plasmids used in this study.

| Name | Description |
| --- | --- |
| pSM2 | pRS316; *TLC1*; CEN; *URA3* |
| pBB39 | pRS426; *pGPD-RNH1-HA*; 2μ; *URA3* |
| pBL97 | pRS316; CEN; *URA3* |
| pBL211 | pRS425; *pGal*; 2μ; *LEU2* |
| pBL352 | pRS425; *pGal-RNH1-HA*; 2μ; *LEU2* |
| pBL189 | pRS426; *pGPD*; 2μ; *URA3* |
| pBL401 | pRS316; *RNH201*; CEN; *URA3* |

### Chromatin Immunoprecipitation

Exponentially grown cells were crosslinked with 1.2% formaldehyde for different time periods. For Sir2 and Rat1-TAP ChIP, cells were crosslinked for 10 min, whereas for pS2-RNA pol II, RNA pol II and Rnh201-TAP ChIP the duration of crosslinking was extended to 15 min. Quenching occurred by incubation for 5 min with 355 mM glycine, followed by cooling on ice for at least 10 min. After 2X washing with 1X cold PBS, the pellets were resuspended in 400 μl cold FA lysis buffer (50 mM HEPES-KOH pH 7.5, 140 mM NaCl, 1 mM EDTA pH 8, 1% Triton X-100, protease inhibitor cocktail) and lysed in lysing Matrix C tubes by FastPrep (MP Biomedicals; 6.5 M/s, 24×2, 3x 30 sec with 1 min on ice between runs). Extracts were recovered with 800 μl cold FA lysis buffer supplemented with 0.1% sodium deoxycholate (FA +SOD) + protease inhibitor cocktail and pelleted. After resuspension with 1.5 ml cold FA +SOD + protease inhibitor cocktail and 0.263% SDS, the cell extracts were sonicated via Bioruptor Pico (Diagenode) at 4°C in order to obtain <500 bp-long fragments. Centrifugation followed and the amount of protein within the supernatant was determined by Bradford. The pellet was discarded and the supernatant employed for further processing as ChIP extract.

A volume of the ChIP extract corresponding to 1 mg/ml protein was combined with cold FA +SOD + protease inhibitor cocktail to obtain a final volume of 1 ml. 50 μl of the resulting mixture was stored at −20°C as the Input. The ChIP extract was incubated with specific amounts of antibodies in combination with the appropriate protein sepharose beads. In details, 5 μg of antibody against RNA pol II and pS2-RNA pol II (Abcam) were used in association with nPprotein A Sepharose 4 Fast Flow beads (GE Healthcare) whereas 10 μg of anti-Sir2 antibody (Santa Cruz) were applied along with Protein G Sepharose 4 Fast Flow beads (GE Healthcare). All ChIP extracts were pre-cleaned for 1 hr with 30 μl beads on a rotor at 4°C in absence of the antibody. After removal of the beads, the antibody was added and the extract was incubated for 1 hr on a rotor at 4°C. Followed the introduction of 50 μl sepharose beads and incubation o/n on a rotor at 4°C. As for TAP-tagged proteins, the pre-cleaning step was omitted and the ChIP extract directly incubated o/n on a rotor at 4°C with 50 μl IgG Sepharose 6 Fast Flow beads (GE Healthcare).

Before usage, all beads were washed with 1X cold PBS, coated with 5% BSA for 1 hr at 4°C and re-washed in 1X cold PBS and cold FA +SOD.

After incubation with the antibody, the ChIP extract was washed with 1 ml cold FA +SOD, FA lysis buffer 500 (FA lysis buffer with 500 mM NaCl), buffer III (10 mM Tris-HCl, pH 8.0, 1 mM EDTA, pH 8.0, 250 mM LiCl, 1% Nonidet P-40 and 1% sodium deoxycholate) and 1X TE (pH 8.0). Afterwards, the IP bound on the washed beads was eluted twice in 100 μl elution buffer (50 mM Tris-HCl, pH 7.5, 1% SDS, 10 mM EDTA, pH 8.0) by incubation for 8 min at 65°C, 1000 rpm. Reverse-crosslinking took place by adding 7.5 μl Proteinase K (0.75 mg/ml; QIAGEN) to the eluate which was then incubated o/n at 65°C. The Input was mixed with 150 μl elution buffer + 7.5 μl Proteinase K and incubated under the same conditions as for the IP. Both the IP and Input were purified with the PCR purification kit (QIAGEN) and eluted in 50 μl elution buffer. qPCR followed by using the CFX384 Touch Real-Rime PCR Detection System (Bio-Rad) and SYBR-Green (Thermo Scientific) (see below for more details about the reaction). The IP values were normalized by corrected Input values, divided by the “% input” values of wild-type cells and the resulting “fold change to wild-types” was represented by GraphPad (Prism7). All oligos used are listed in Table 3.

**Table 3:**
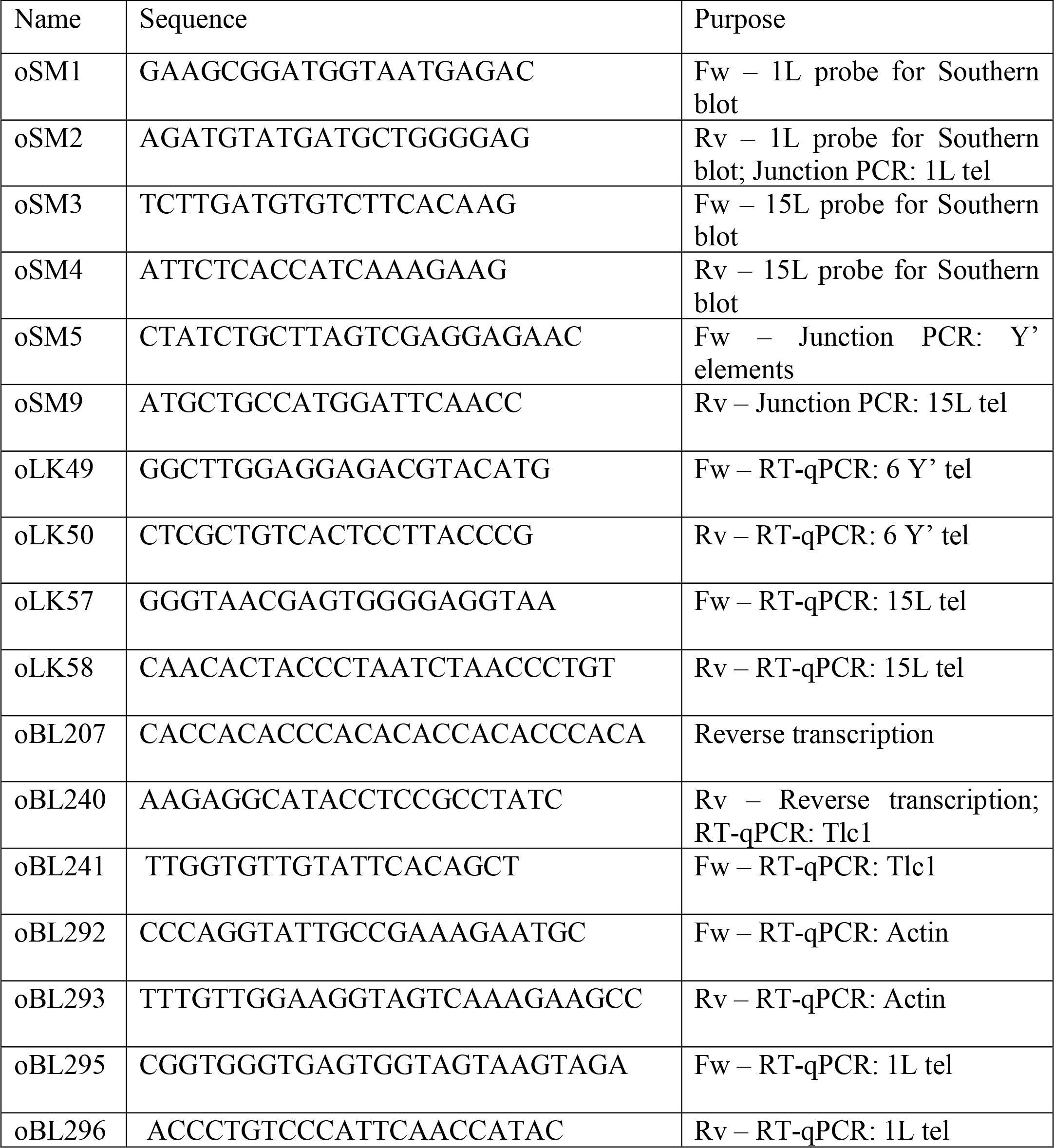
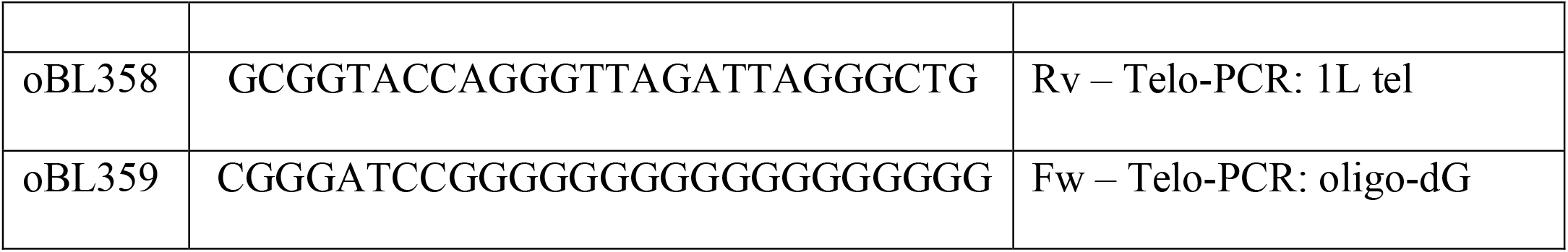
Oligonucleotides used in this study.

### DNA extraction

Exponentially grown cells were pelleted and resuspended in 1 ml of a solution containing 0.9M sorbitol + 0.1M EDTA pH 8.0. After pelleting, cells were mixed with 400 μl of the same solution supplemented with 14 mM β-mercaptoethanol and incubated with 20 μl of 2.5 mg/ml Lyticase for 1 hr at 37°C. After checking the formation of spheroblasts, cells were pelleted, resuspended in 400 μl 1X TE and mixed with 90 μl of a solution containing 0.3 M EDTA pH 8.0, 0.2 M Tris-Base and 2.2% SDS. Followed an incubation for 30 min at 65°C, at the end of which 80 μl of 5 M KAc were added. The samples were moved directly on ice for at least 1 hr and then centrifuged at 4°C for 15 min at maximum speed. The supernatant was transferred into 750 μl cold 100% EtOH and incubated for 30 min −20°C, after gentle mixing. The samples were then centrifuged for 5 min at 4°C and the precipitate was washed with 1 ml 70% EtOH and left air-drying. Once dry, the precipitate was resuspended in 500 μl 1X TE followed by RNA digestion with 2.5 μl RNase A (Thermo Scientific) for 30 min at 37°C. At this point, 500 μl Isopropanol were added and, after gentle mixing, the samples were incubated for 30 min at −20°C. Centrifugation for 15 min at 4°C took place and the precipitate was washed with 1 ml 70% EtOH before air-drying. The precipitate was then eluted in 50 μl 1X TE. Once fully resuspended, the eluate was centrifuged for 10 min at 4°C and the supernatant stored at −20°C.

### Southern blot

Approximately 5 μg genomic DNA were digested with 1.5 μl restriction enzymes + 2.5 μl CutSmart buffer (NEB) for 5 hrs at 37°C. The restriction enzyme XhoI (NEB) was used when detecting all telomeres and telomere 15L, whereas SalI (NEB) was employed to study telomere 1L. At the end of digestion, the samples were mixed with 1X loading dye and loaded on a 0.8% agarose gel. Electrophoresis took place o/n at 50-60 V. After migration, the DNA in the gel was first denatured with 0.4 M NaOH, 0.6 M NaCl for 1 hr and then neutralized in 1 M Trizma Base, 1.5 M NaCl (pH 7.4) for another hour. DNA was transferred onto a nylon membrane (Hybond-NX, GE Healthcare) via capillarity in 10X SSC over 72 hrs. After the transfer, the DNA was crosslinked with UV light (Auto X-link, Stratalinker) on the membrane, which was then pre-hybridized with PerfectHyb buffer (Sigma-Aldrich) o/n at 55°C. Hybridization with radioactive telomeres-specific probes followed o/n at 55°C. To generate the CA-probe, which recognizes the entire telomere population, a ∼ 300 bp-long plasmid-derived fragment containing a telomeric sequence was used as a template for random labeling (DECAprime II DNA Labeling Kit, Thermo Scientific) with dATP [α-^32^P] (Perkin Elmer/Hartmann-Analytic). Differently, the substrate for the probe detecting telomere 1L and 15L was derived from a PCR reaction, using primers specific for the subtelomeric region (Craven and Petes, 1999). After hybridization with the radioactive probes, the membrane was washed twice first with pre-warmed 2X SSC, 0.1% SDS and then with 0.5X SSC, 0.1% SDS (5 and 20 min, respectively) at 55°C. The membrane was left to air-dry for approximately 1 hr and exposed to a phosphorimaging film for 3-4 nights. Detection of the signal was carried out via Typhoon FLA 9500 (GE Healthcare).

### Telomere length analysis

Telomere length was determined by using Image J. Images of Southern blots were rotated 90° to the right in order to have them perpendicular to the x-axis. The left edge of the picture corresponded to x=0 and the telomere length was defined as the distance of each telomeric band on the blot from x=0 on the x-axis. Virtual lanes containing each band were generated by the Analyze tool. After setting the first lane (Analyze > Gels > Select First Lane), the subsequent lanes were formed by the command Analyze > Gels > Select Next Lane. Once defined, all lanes were plotted with the command Analyze > Gels > Plot Lanes and each peak corresponding to the most intense area of the bands selected in the profile with the Multi-point tool. At the end, by using the command Analyze > Measure, the distance of each band from x=0 was calculated and reported as an x value. The x value corresponding to the longest telomere was set as 1 and the other x values were divided by it to obtain the relative loss of telomere length.

### Calculation of the telomere shortening rate

Two approaches were used. In Figure 5A, the length of telomere 1L measured by Telomere-PCR was subtracted from the one corresponding to the previous time point. The resulting value was divided by the difference between the population doublings of the two time points to obtain the final “bp/PD”.

In Figure 5B, the slope was derived from the shortening profile of telomere 1L on the left part of the panel for each replicate by using the “simple linear regression” function of GraphPad (Prism).

### Telomere-PCR

1 μl containing 100 ng genomic DNA, extracted as previously annotated, was mixed with 7.1 μl H_2_O and 0.9 μl NEB 4 buffer (NEB). After incubating for 10 min at 96°C, the mix was cooled down to 4°C and supplemented with 1 μl of a solution containing 4 U Terminal Transferase (Thermo Scientific), 0.1 μl 10X NEB 4 buffer (NEB), 0.1 μl 10 mM dCTP and H_2_O. The samples were then incubated for 30 min at 37°C, 10 min at 65°C and 5 min at 96°C. Afterwards, the samples were kept at a temperature of 65°C. Meanwhile, 21 μl H_2_O were mixed with 4 μl 10X PCR buffer (89.11 mM Tris-HCl pH 8.8, 21.28 mM (NH4)2SO4, 6.65% glycerol and 0.0133% Tween-20), 4 μl dNTPs (2 mM stock), 0.3 μl forward subtelomeric oligo (100 μM stock), 0.3 μl G18 reverse oligo oBL359 (100 μM stock) and heated at 65°C. At this temperature, 0.5 μl Phusion Hot Start II DNA Polymerase (Thermo Scientific) were added and the resulting mix was combined with the samples at 65°C. Thermocycling took place as follows: 3 min at 95°C, 45 cycles of 30 sec at 95°C, 15 sec at 63°C and 20 sec at 68°C. At the end, the samples were held for 5 min at 68°C and then at 4°C. The derived PCR products were separated on a 1.8% agarose gel for 2.5 hrs at 100V. The telomere length was defined by Image Lab or Image J.

### RNA extraction

Cells from overnight cultures were exponentially grown (OD600: 0.6-0.8) in 15 ml medium and pelleted. Pellet was resuspended in 400 μl AE-buffer (50 mM NaAc pH 5.3, 10 mM EDTA) and combined with 20 μl 20% SDS and 500 μl phenol pre-equilibrated in AE-buffer. The resulting mix was heated for 5 min at 65°C followed by incubation on ice for 5 min. Subsequently, the samples were centrifuged at 14000 rpm for 3 min at 4°C and the supernatant was mixed with 500 μl phenol-chloroform (1:1). The mix was Incubated for 5 min at room temperature and centrifuged as before. The supernatant was mixed with 40 μl 3M NaAc and 1 ml 100% ethanol and incubated for 30 min at −20°C. Afterwards, the samples were centrifuged as before and the pellet was washed with 1 ml 80% ethanol. To remove DNA, the pellet was resuspended in a solution composed of 86 μl H_2_O, 10 μl RDD buffer (QIAGEN) and 4 μl DNase I (QIAGEN) and incubated for 1 hr at 37°C. 50 μg RNA were further purified by performing three times the clean-up procedure with the RNeasy kit (QIAGEN). DNA digestion was carried out as before between each clean-up step. During the last clean-up, RNA was eluted in 30 μl H_2_O.

### Reverse transcription and RT-qPCR

3 μg purified RNA were diluted in H_2_O to obtain a final volume of 7 μl. Subsequently, RNA was combined with 0.4 μl dNTPs (25 mM each stock), 1 μl oligo oBL207 (10 uM stock), 0.4 μl oligo oBL293 (10 uM stock) and 4.2 μl H_2_O. The resulting mix was heated for 1 min at 90°C and cooled down to a temperature of 55°C with a rate of about 0.8°C/sec. At this temperature, a mix composed of 1 μl DTT (0.1 M stock, Invitrogen), 1 μl SuperScript III Reverse Transcriptase (Invitrogen), 1 μl RNase OUT (Invitrogen) and 4 μl First Strand buffer (5X stock, Invitrogen) was added to the samples. cDNA synthesis was carried out for 1 hr at 55/65°C, followed by heat inactivation of the enzyme for 15 min at 70°C. As a negative control, RNA samples were subjected to the same process but in the presence of H_2_O instead of the reverse transcriptase. 30 μl H_2_O were added at the end.

For the RT-qPCR, 1 μl cDNA was mixed with 1 μl each primer, 2 μl H_2_O, 5 μl SYBR-Green (Thermo Scientific) and the reaction was performed on the CFX384 Touch Real-Time PCR Detection System (Bio-Rad) by denaturing DNA for 10 min at 95°C, followed by 40 cycles of 15 sec at 95°C and 1 min at 60°C to quantify the signal. TERRA levels were normalized to the actin mRNA levels by calculating ΔCt =Ct TERRA-Ct actin.

### Protein extraction and Western blot

2 OD600 units of exponentially growing cells were pelleted, resuspended in 150 μl of a solution containing 1.85 M NaOH, 1.09 M β-mercaptoethanol and incubated for 10 min on ice. Afterwards, the samples were mixed with 150 μl of a 50% TCA solution and incubated for additional 10 min on ice. Centrifugation at 4°C took place and the pellets were resuspended in 1 ml acetone. After another centrifugation at 4°C, the pellets were resuspended in 100 μl urea buffer (120 mM Tris-HCl pH 6.8, 5% glycerol, 8 M urea, 143 mM β-mercaptoethanol, 8% SDS, bromophenol blue indicator) and stored at −80°C.

For Western blotting, samples were thawed at room temperature before being heated for 5 min at 75°C. 8 μl from each sample were loaded onto Mini Protean TGX Pre-cast 4-15% gels (Bio-Rad) and migration was carried out for approximately 45 min at 150V. Gels were blotted on nitrocellulose membranes and transfer was carried out by using the “high molecular weight” program of the Trans-Blot Turbo System (Bio-Rad). The membranes were blocked in a blocking solution (5% skim milk, 1X PBS, 0.1% Tween-20) for 1 hr at room temperature and incubated o/n at 4°C with the appropriate primary antibodies diluted in blocking solution. Afterwards, the membranes were washed four times for 15 min at room temperature with a washing solution (1X PBS, 0.1% Tween-20) and, when required, incubated for 1 hr at room temperature with the proper HRP-coupled secondary antibodies diluted in blocking solution. The membranes were washed anew three times for 15 min with the washing solution and incubated for 1 min with Super Signal Pico or Dura Chemiluminescent Substrate (Thermo Scientific). Detection was performed with the “Chemiluminescence” program of the ChemiDoc Touch Imaging System (Bio-Rad).

### Replicative potential measurement

Survivors were streaked for single colonies on solid selective medium or YPD. When colonies formed, they were individually inoculated in liquid selective medium or YPD and grown o/n at 30°C. The following day, the OD600 was measured and cells were passed into fresh medium to obtain a final OD600: 0.01 and grown o/n at 30°C. The OD600 measurement, dilution to OD600: 0.01 and o/n growth were carried out every 24 hrs.

The replicative potential was determined by setting the first OD600 value of each survivor clone transformed with an empty vector (EV, Figure 4A) or with the Rnh201-expressing plasmid (final genotype: *tel-*, Figure 4B) as 1 and by dividing the OD600 values obtained for the same clone transformed with an EV or the RNH1-overexpressing/Rnh201-expressing plasmid on the same day and the following days by it. The resulting value indicates the proliferative ability that cells have in relation to the initial replicative potential.

When survivors were monitored for TERRA levels during the clonal propagation, cells were isolated from the overnight cultures on different days and exponentially grown. After collecting the pellet, the procedure to examine TERRA was the same as described above.

### Survivor associated senescence rate measurement

The senescence rate in survivors was defined as the number of population doublings (PDs) required for telomeres to reach the shortest length starting from the beginning of the experiment (“Start” in Figure 4E and F). This was assumed to coincide with the lowest value of replicative potential, or local minimum, during the clonal propagation (“Min” in Figure 4E and F). In order to avoid biases given by stochastic fluctuations of the OD600 measurement, the “% replicative potential” profiles were smoothened by calculating the average of two consecutive values. Among these values, local minima were considered only if no other values in a neighborhood of four were inferior. The similarity between the proliferative potential profiles of the conditions tested, e.g., EV vs RNH1, was also used as a qualitative confirmation that the correct minima were compared.

## Results and Discussion

### TERRA regulators are less abundant at type II survivors telomeres

Increased TERRA levels is a hallmark of human ALT cancer cells and plays a role in HDR, which drives telomere maintenance in telomerase-negative cancers^31,32,34,36–39^. Consistently, in both budding and fission yeast, post-senescent survivors, which also use HDR for telomere extension, present high levels of TERRA^35,40^. The origin of TERRA upregulation remains to be fully clarified. In humans, increased transcription seems to be the primary source of increased TERRA as demonstrated by the enhanced telomeric enrichment of RNA pol II and the similar turnover rates of TERRA in ALT cells compared to telomerase-positive cells^31,34^. ALT-associated hypomethylation of TERRA promoters and reduced chromatin condensation at telomeres are proposed to fuel the process^31,34^.

To understand the mechanisms underlying TERRA accumulation in type II survivors in budding yeast, we monitored the telomeric association of previously characterized TERRA regulators through chromatin immunoprecipitation (ChIP). Sir2, the catalytic subunit of a histone deacetylase complex that negatively regulates TERRA, is less enriched at telomeres 1L, 15L and 6 Y’ telomeres in survivors compared to telomerase-positive wild-type cells (Figure 1A). The decreased association was restricted to telomeres, as Sir2 ChIP at the mating locus HML was unaffected by the survivor status of the cells. These results indicate that, similar to human ALT cells, telomeric chromatin may be altered in type II survivors. To test whether transcription was increased as a result, the recruitment of RNA pol II, both total (RNAPII) and the elongating form (pS2-RNAPII), was assayed at telomeres. No differences were observed between survivors and wild-type cells, suggesting that TERRA upregulation in survivors may not be due to increased transcription despite the lack of Sir2 associated to chromatin (Figure 1B and Supplementary Figure S1A).

**Figure 1.**
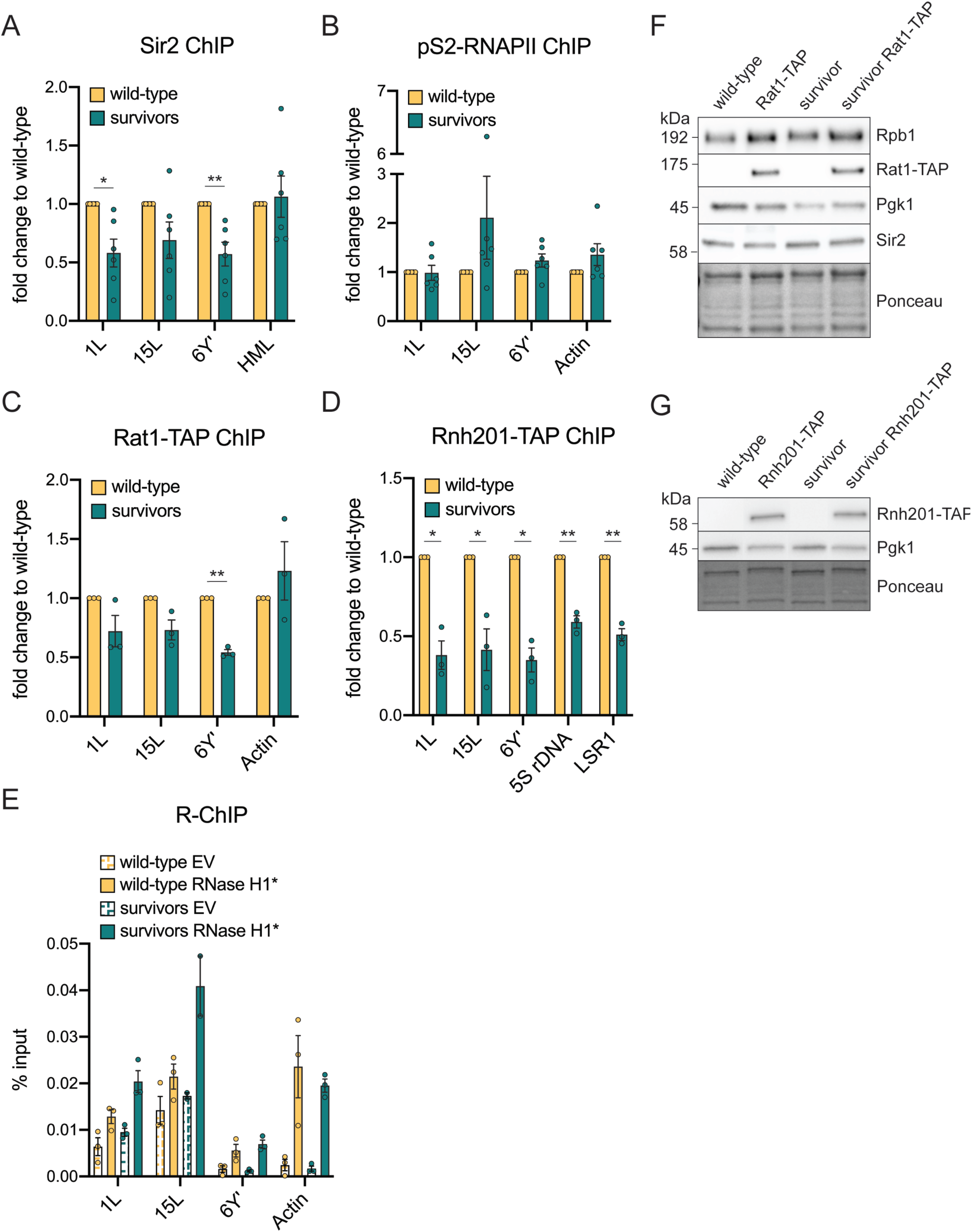
TERRA regulators are reduced at survivor telomeres. **(A)** Sir2 ChIP was performed in *tlc1Δ* type II survivors and wild-type cells at three telomeric loci (1L, 15L and 6Y’) and the mating locus HML, used as positive control. The “% input” value of each survivor was divided by the “% input” of the isogenic wild-type collected on the same day to obtain the final “fold change to wild-type”. **(B)** ChIP for the elongating version of RNA pol II (phosphorylated at Ser2) was performed as described in A) with the exception that the actin locus was used as a positive control. **(C)** ChIP for the TAP-tagged Rat1 was performed in isolates from *tlc1Δ* survivors and wild-type cells harboring the tagged and untagged version of the protein. The above indicated telomeres were monitored for the protein enrichment along with the actin locus, used as positive control. The “% input” values of TAP-tagged survivors and wild-type cells were divided by the ones of the untagged equivalents and the resulting fold change was divided by the value of wild-type cells to obtain the final “fold change to wild-type”. **(D)** ChIP was performed for TAP-tagged Rnh201 in *est2Δ* survivors and wild-type cells. The 5S rDNA and LSR1 loci were used as positive controls. The “fold change to wild-type” value was calculated as in C). **(E)** R-loop levels at 1L, 15L and 6Y’ telomeres in wild-type cells and *tlc1Δ* survivors measured by R-ChIP. Cells were transformed with an empty vector (EV) or a vector expressing the catalytically dead version of RNase H1 (RNase H1*). Expression was induced with galactose for 2 hours. The protein enrichment at the indicated loci is normalized by the input. The actin locus was used as positive control. **(F)** Western blot for Rpb1, Rat1-TAP and Sir2 in *tlc1Δ* survivors and wild-type cells with and without the TAP tag on the Rat1 protein. **(G)** Western blot for Rnh201-TAP in survivors and wild-type cells with or without the tag on the Rnh201 protein. Pgk1 detection and Ponceau staining were used as loading controls. In (A) and (B), 6 replicates were used (n=6) while in C and D, 3 replicates were used (n=3). Mean + SEM is displayed and p values were calculated by an unpaired two-tailed Student’s *t*-test with Welch’s correction (*p<0.05, **p<0.01).

TERRA turnover is primarily mediated by the nuclease activity of Rat1, which is recruited to yeast telomeres, in part, via the Rif proteins^27,29^. To investigate whether Rat1 mis-localization may account for TERRA upregulation in survivors, as it does in pre-senescent cells, we performed ChIP on Rat1-TAP-expressing cells^29^. Rat1 was less enriched at the chromosome ends of survivors, especially at Y’ telomeres, where binding was reduced by almost 50% (Figure 1C). This suggests that survivor-associated accumulation of TERRA may be in part due to less Rat1 at telomeres, and hence reduced turnover.

TERRA forms RNA-DNA hybrids (presumably R-loops) at telomeres in both humans and yeast^28,31^. In human ALT cancer cells, R-loop accumulation causes replicative stress that triggers HDR at telomeres^31,33^. Consistently, in budding and fission yeast, RNA-DNA hybrid removal impairs survivor formation and proliferation^35,40,41^. We performed ChIP for Rnh201, the catalytic subunit of the RNase H2 enzyme, which participates in the removal of RNA-DNA hybrids at telomeres^29^. In survivors, Rnh201 association to telomeres was strongly reduced as compared to wild type cells (Figure 1D). The reduced chromatin association was not telomere specific as shown by the lower enrichment also at other loci. This indicates that type II survivors might accumulate RNA-DNA hybrids not only at the chromosome ends but also at other sites in the genome. To verify that RNA-DNA hybrids were indeed increased at telomeres in survivors, we employed ChIP experiments with catalytically dead RNase H1 (termed R-ChIP). Consistent with loss of RNase H2 at telomeres, we detected an increased R-ChIP signal at telomeres in survivors as compared to wild type cells (Figure 1E). Importantly, the total protein amount of Sir2, RNA pol II, Rat1 and Rnh201 was unchanged in survivors compared to telomerase-positive cells, thus excluding differences in expression as the source of unequal binding of the proteins to telomeres (Figure 1F, G). Taken together, these data indicate that survivors present a more open chromatin state with less RNase H2 and Rat1 association. Overall, this suggests that TERRA abundance may be enhanced due to decreased turnover rates through RNase H2 and Rat1, rather than increased transcription.

### TERRA regulation throughout the cell cycle is not altered in type II survivors

TERRA expression is modulated throughout the cell cycle in humans and yeast, being upregulated at the beginning of S phase and declining as replication proceeds and cells enter G2 phase^29,42–44^. In human ALT cancer cells, this regulation may be altered, as TERRA’s telomere association remains when cells enter the G2 phase^43^.

To test whether TERRA upregulation in yeast survivors resulted from perturbations of its cell cycle regulation, TERRA expression was monitored in type II survivors in different cell cycle phases and compared to wild-type cells as previously described^29^. α-factor was used to arrest the cells in G1 whereas 200 mM and 75 mM hydroxyurea were employed to block the cells in early and late S phase, respectively. In line with previous results, TERRA levels were very low in G1, increased in early S phase and diminished by approximately 29% as replication completed and cells entered the G2 phase (Supplementary Figure S1B)^29^. Survivors displayed the same cell cycle regulation, with TERRA peaking in early S phase and dropping by approximately 33.5% at the end of replication. Although the cell cycle regulation was similar, the amount of TERRA from telomeres 1L and 15L was significantly higher in survivors than wild-type cells in each cell cycle phase, consistent with it being upregulated in survivors^35^. In wild-type cells, the cell cycle regulation (from early S to G2) is dependent on Rat1-mediated degradation: TERRA is transcribed in S phase and Rat1 degrades it in G2 and G1^29^. Therefore, despite the slightly reduced Rat1 binding to telomeres in survivors (Figure 1C) it appears to be sufficient to maintain the late S/G2 degradation. We speculate that in the case of survivors, it may be the loss of Sir2 that is largely responsible for increased TERRA levels. This would better reflect the scenario described in human ALT cells, where telomeres are more transcribed due to less chromatin compaction and the rates of TERRA turnover are not altered^34^. It is also possible that the regulation of TERRA is only lost at a small subset of telomeres in survivors. Indeed, telomeres in type II survivors are extremely heterogeneous in length and may differ in the way they regulate the lncRNA expression.

### Telomerase expression reduces TERRA levels in survivors

We have previously shown that senescent cells accumulate TERRA at short telomeres due to the reduced binding of Rat1^29^. Whereas type II survivors are typically described as having long and heterogeneous telomeres, it is often overlooked that they also accumulate many critically short telomeres (Figure 2A, black arrow). Since survivors present critically short telomeres and, similar to senescent cells, have increased TERRA levels, we hypothesized that the pool of critically short telomeres in type II survivors may be responsible for the increase of TERRA as is the case in senescent cells. To address this, we re-introduced telomerase into established type II survivors through transformation of *TLC1*, the RNA component of telomerase, and allowed cells to undergo at least 30 population doublings. Telomerase expression was similar in wild-type cells as compared to type II survivors (Supplementary Figure S2A). As expected, telomerase preferentially elongated the critically short telomeres, while the overall structure and heterogeneity of type II survivors telomeres was maintained (Figure 2A)^45^. The presence of extra copies of *TLC1* did not alter the telomere length in wild-type cells (Figure 2A).

**Figure 2.**
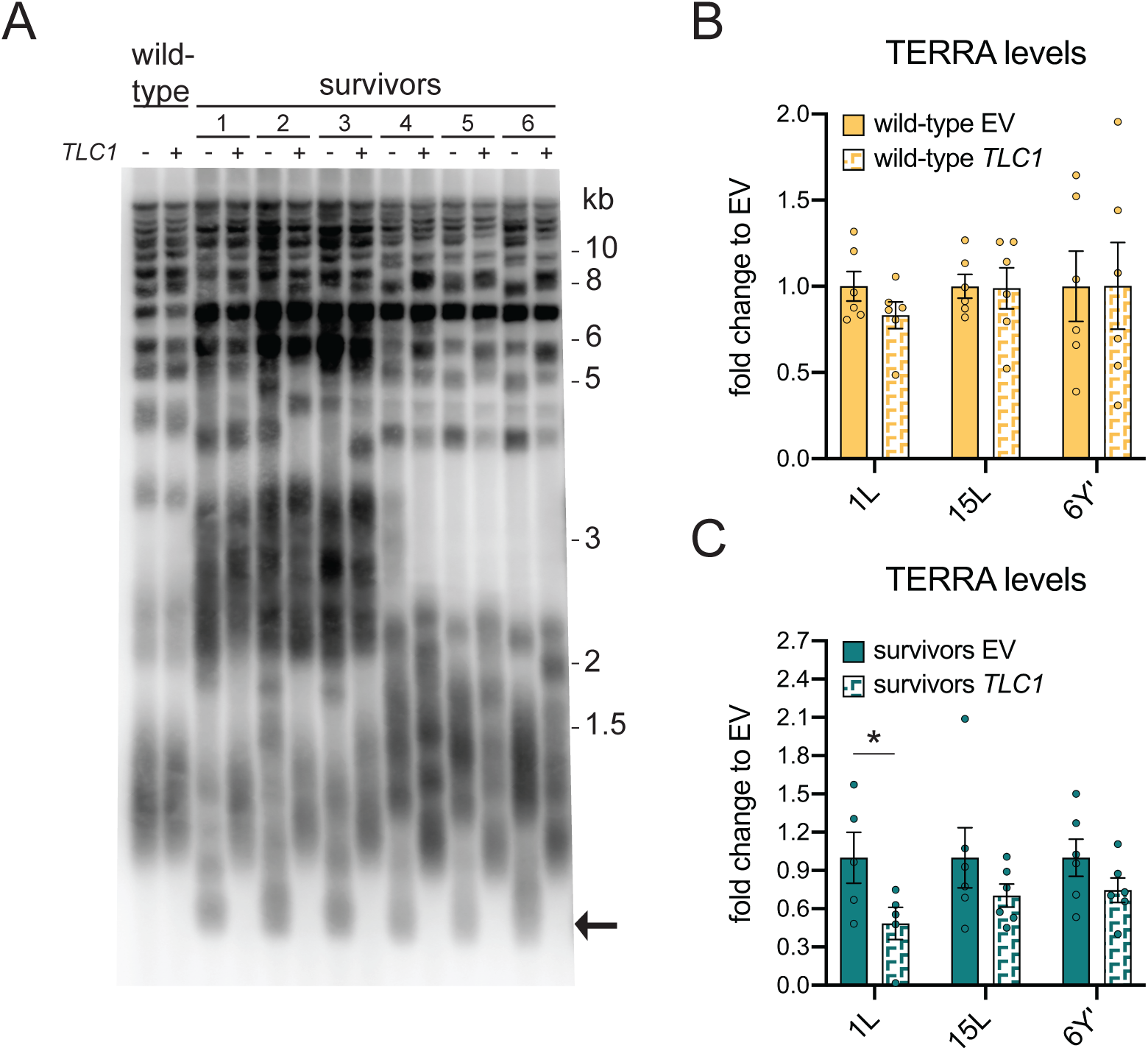
Telomerase expression reduced TERRA upregulation in survivors. **(A)** Southern blot for all telomeres in six *tlc1Δ* survivor clones and one wild-type clone transformed with a vector carrying the *TLC1* gene under its endogenous promoter (+) or an empty vector (-). The arrow points to the critically short telomeres that exist in type II survivors. **(B)** TERRA levels were measured at 1L, 15L and 6Y’ telomeres in wild-type cells (one of the clones is shown in A) transformed with the *TLC1*-carrying vector (*TLC1*) or an empty vector (EV). The amount of TERRA of each sample was determined as “% actin mRNA” and divided by the average among the wild-type cells transformed with the EV to obtain the final “fold change to EV”. **(C)** TERRA levels were measured at 1L, 15L and 6Y’ telomeres in *tlc1Δ* type II survivors from A), as described in B). The amount of TERRA of each sample was determined as “% actin mRNA” and divided by the average among the survivors transformed with the EV to obtain the final “fold change to EV”. Mean + SEM is displayed, n=5/6. p values were calculated by an unpaired two-tailed Student’s *t*-test (*p<0.05).

TERRA levels were unaffected by the introduction of exogenous *TLC1* in wild-type cells (Figure 2B). In contrast, telomerase expression in survivors caused a reduction of TERRA expression between 30%-50% (Figure 2C). These data suggest that short telomeres in type II survivors may be contributing to the increased levels of the lncRNA. Importantly, a similar effect was observed in human ALT cancer cells, where overexpression of telomerase caused a decrease of TERRA expression. These authors also suggested that increased TERRA in human ALT cells is stemming from a pool of short telomeres, as the re-introduction of telomerase also elongates short telomeres in ALT cancer cells^34,46,47^. Furthermore, it was demonstrated that in human cells, the re-introduction of telomerase into ALT cells increases chromatin compaction which they attribute to being responsible for the reduced TERRA levels^34^. This is consistent with the decreased Sir2 association detected at survivor telomeres (Figure 1A). The reduction in TERRA levels is quite subtle and levels varied considerably between clones. This is likely due to survivor heterogeneity combined with the fact that only a small subset of telomeres is critically short. Hence, to corroborate these results it will be necessary to follow telomere shortening in individual survivor clones (see below).

### Type II survivors regulate TERRA in a telomere length-dependent manner

So far, type II survivors had been generated by allowing clonal telomerase-negative cells to enter crisis following serial passage in a liquid culture (Figure 3A). The recovery of the culture then becomes populated with type II survivors as confirmed by Southern blotting. Using this method, the culture maintains a high replicative potential and single telomere length fluctuations are not detectable. However, such a type II culture is highly heterogeneous as multiple cells escape senescence and the type II culture does not arise from a single cell. To better understand how TERRA and telomere length are regulated in survivors, we isolated single clones from independently generated type II survivors (Figure 3A) as well as from wild-type cultures and propagated them in liquid medium for almost 800 population doublings; samples for DNA and RNA samples were taken throughout the experiment in order to measure telomere length and TERRA levels, respectively (Figure 3A). By using individual clones instead of mixed populations of cells we could follow the dynamics of single telomeres with unique lengths rather than a pool with various lengths. We studied the length variation of telomere 1L and 15L by Southern blots in which the probe recognized the specific subtelomeric regions, as described by Craven and Petes (Figure 3B)^48^.

**Figure 3.**
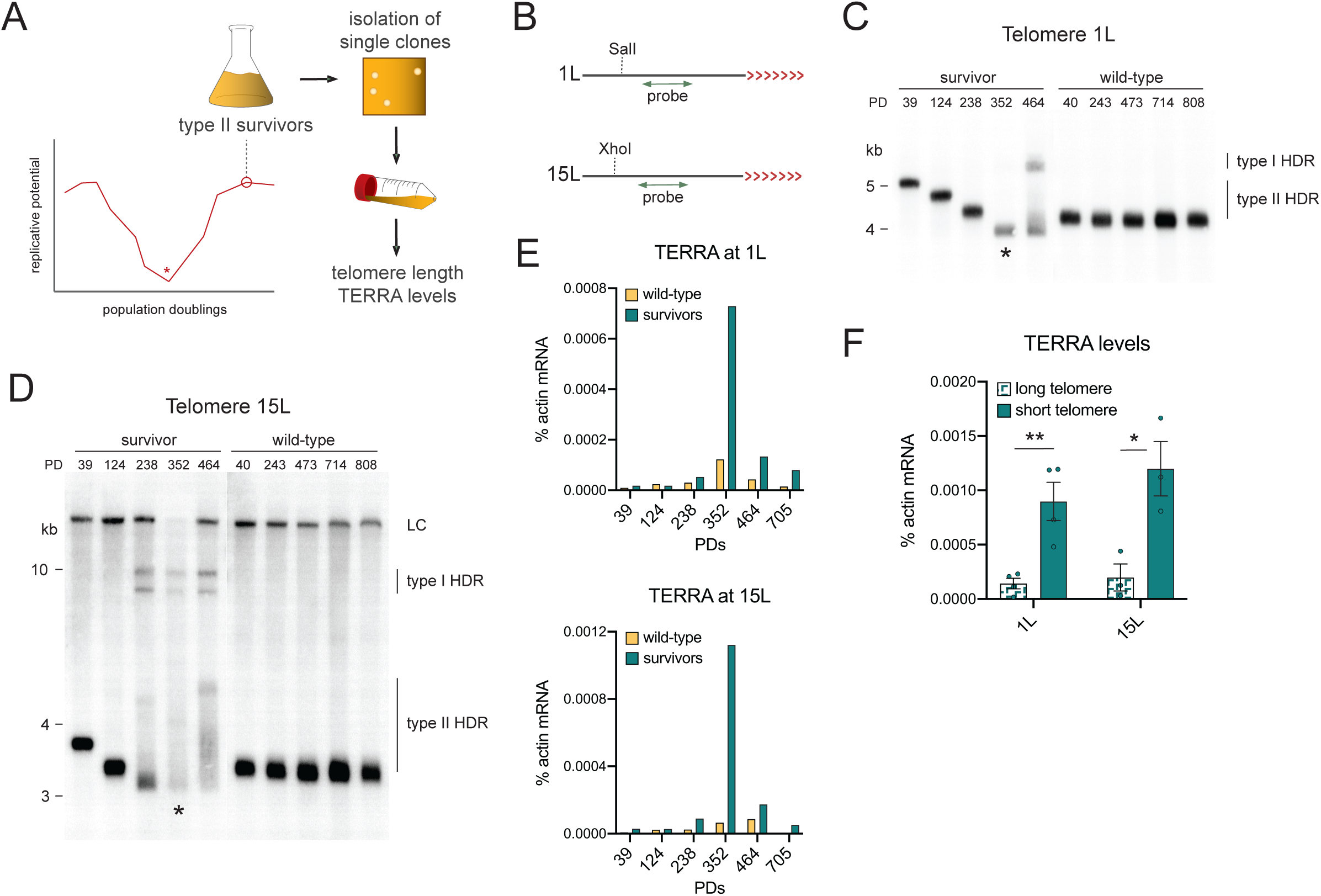
TERRA levels increase as telomeres shorten in survivors. **(A)** Schematic of the experimental setup to generate single clones of type II survivors. Briefly, individual clones of independently generated *tlc1Δ* survivors were isolated on solid medium and propagated in liquid. DNA and RNA were extracted from the same monoclonal population at specific time points during propagation to measure the telomere length and the TERRA levels, respectively. The asterisk indicates the time point when senescent cells enter crisis before giving rise to survivors. **(B)** Schematic of the subtelomere/telomere structure at the 1L/15L chromosome ends. SalI and XhoI are the restriction enzymes used to digest the genomic DNA in preparation of telomere 1L and 15L Southern blotting, respectively. The red arrowheads represent the telomere whereas the black line indicates the subtelomere. The double arrowheads in green represent the radioactive probes used to detect the single telomeres. **(C)** Southern blot for telomere 1L in one *tlc1Δ* survivor and wild-type clone at different population doublings (PD) during the monoclonal propagation. The asterisks indicate the time point at which the population presented the shortest telomere length. **(D)** Southern blot for telomere 15L in the same samples used for (C). Bands recognized by non-specific binding of the 15L-probe were marked as loading control (LC). In both Southern blots (C and D), the type of HDR is indicated. **(E)** TERRA levels measured as “% actin mRNA” in the same samples used in C) and D). **(F)** The amount of TERRA generated from the longest telomere 1L and 15L was compared to the one from the respective shortest telomere in the survivor shown in C) and D) as well as from additional independently generated *tlc1Δ* survivors (Supplementary Figure S3A-C). Mean + SEM is displayed, n=4. p values were calculated by an unpaired two-tailed Student’s *t*-test (*p<0.05, **p<0.01).

Surprisingly, telomeres 1L and 15L gradually lost their repeat length (Figure 3C, D), similar to what has been reported for telomerase-negative pre-senescent cells and type II survivors harboring a modified telomere^22,49,50^. When telomeres became critically short (i.e., less than wild-type length), telomeres were re-extended, as indicated by the appearance of longer restriction fragments. In particular, type I HDR promoted the acquisition of Y’ elements while type II HDR resulted in the addition of different extents of TG-repeats. Hence, after “crisis” the population of survivors became heterogeneous again with different cells using either a type I or type II HDR. As expected, the length of telomere 1L and 15L changed very little in wild-type cells, consistent with telomerase being active throughout the course of the experiment. To our knowledge, this is the first report that a natural telomere in survivors undergoes shortening in a similar manner to pre-senescent cells. Moreover, these data indicate that in survivors, telomeres are not constantly undergoing HDR but only do so when they become critically short. Therefore, the re-extension of short telomeres in type II survivors by HDR is analogous to telomerase’s preference for short telomeres in telomerase-positive cells^45^.

During replicative senescence, TERRA levels progressively increase as telomere length diminishes, and increases dramatically at critically short telomeres^29^. We observed a similar relationship in type II survivors, with TERRA levels peaking at the population doubling time when critically short telomeres appeared (Figures 3E, F and Supplementary Figures S3A-C). Upon analyzing multiple survivors, we found that on average, in survivor clones with short telomeres (below wild type length), TERRA accumulated to approximately 6 times more than survivors with long telomeres (Figure 3F, S3). When telomeres were re-extended following crisis, the levels of the transcript declined back to pre-crisis levels. These data, together with reduced TERRA levels following the re-expression of telomerase in survivors (Figure 2), strongly suggest that increased TERRA levels in survivors stems from short telomeres. Moreover, these data demonstrate that survivors also go through cycles of telomere shortening and re-elongation in a similar manner to pre-senescent cells, with TERRA levels responding accordingly. Finally, it appears that the telomere length-dependent regulation of TERRA in yeast survivors may be similar to what has been speculated in human ALT cancer cells^34^.

Considering the similarity between the regulation of TERRA in survivors and pre-senescent cells, it is tempting to think that short telomeres in type II survivors accumulate TERRA due to reduced turnover rates as well. Although only a mild reduction of Rat1 was observed at telomeres in survivors compared to wild-type cells (Figure 1C), this may be due to the fact that only a subpopulation of survivors has short telomeres in a mixed population culture. The differences observed for the Sir2 and Rnh201 telomere association may also be due to reduced binding at short telomeres, however, since the reduction is more accentuated for Sir2 and Rnh201 than for Rat1, we cannot exclude that also long telomeres have less Sir2 and Rnh201 in survivors. Type II survivors would therefore resemble their human ALT cells in having a more open chromatin at telomeres regardless of their length^34^.

Overall, we propose a model where TERRA and its association to telomeres increase as telomeres shorten in type II survivors. This is likely to occur because less negative regulators can associate to short telomeres. When telomeres are critically short (below the wild-type length), the high levels of TERRA and perhaps RNA-DNA hybrids may stimulate HDR-mediated extension.

### Type II survivors undergo senescence

Type II survivors appear to have a common phenotype with senescent cells in that their telomeres, although initially very long, progressively shorten and accumulate TERRA (Figure 3)^29^. During replicative senescence, TERRA RNA-DNA hybrids counteract replicative senescence, presumably by either preventing telomere shortening or promoting HDR-mediated extension^28^. To address whether RNA-DNA hybrids have the same “anti-senescence” effect in survivors, we monitored the replicative potential in type II survivors with different amounts of RNA-DNA hybrids. We propagated isolated survivor clones with either reduced (RNase H1 overexpression) or increased (deletion of RNase H2) abundance of telomeric RNA-DNA hybrids and compared their proliferation potential to isogenic cells with unperturbed levels of hybrids, as previously described^28,29^. To examine the effect of decreased R-loops, we transformed type II survivors with an *RNH1*-overexpressing plasmid or an empty vector control (EV) prior to cell passaging. In cells harboring an EV control, we noticed that the replicative potential of most survivors fluctuated as cells were propagated for approximately 250 population doublings, with maximal values being up to 10 times higher than the minimal ones (Figures 4A, B yellow lines). To understand if the variations of replicative potential corresponded to changes in TERRA levels, as they do during replicative senescence, we monitored the TERRA levels of telomere 15L in one of the clones (from Figure 4A) at different time points. Cells of survivor C transformed with an empty vector progressively lost their ability to proliferate from PD 35 to PD 149. TERRA levels increased as cells lost their replicative potential consistent with telomere shortening (Figure 4C). Once lengthening took place and the ability to divide was restored, the transcript levels declined. This is the first report of survivors displaying a senescence phenotype that we refer to as “survivor associated senescence” or “SAS”.

**Figure 4.**
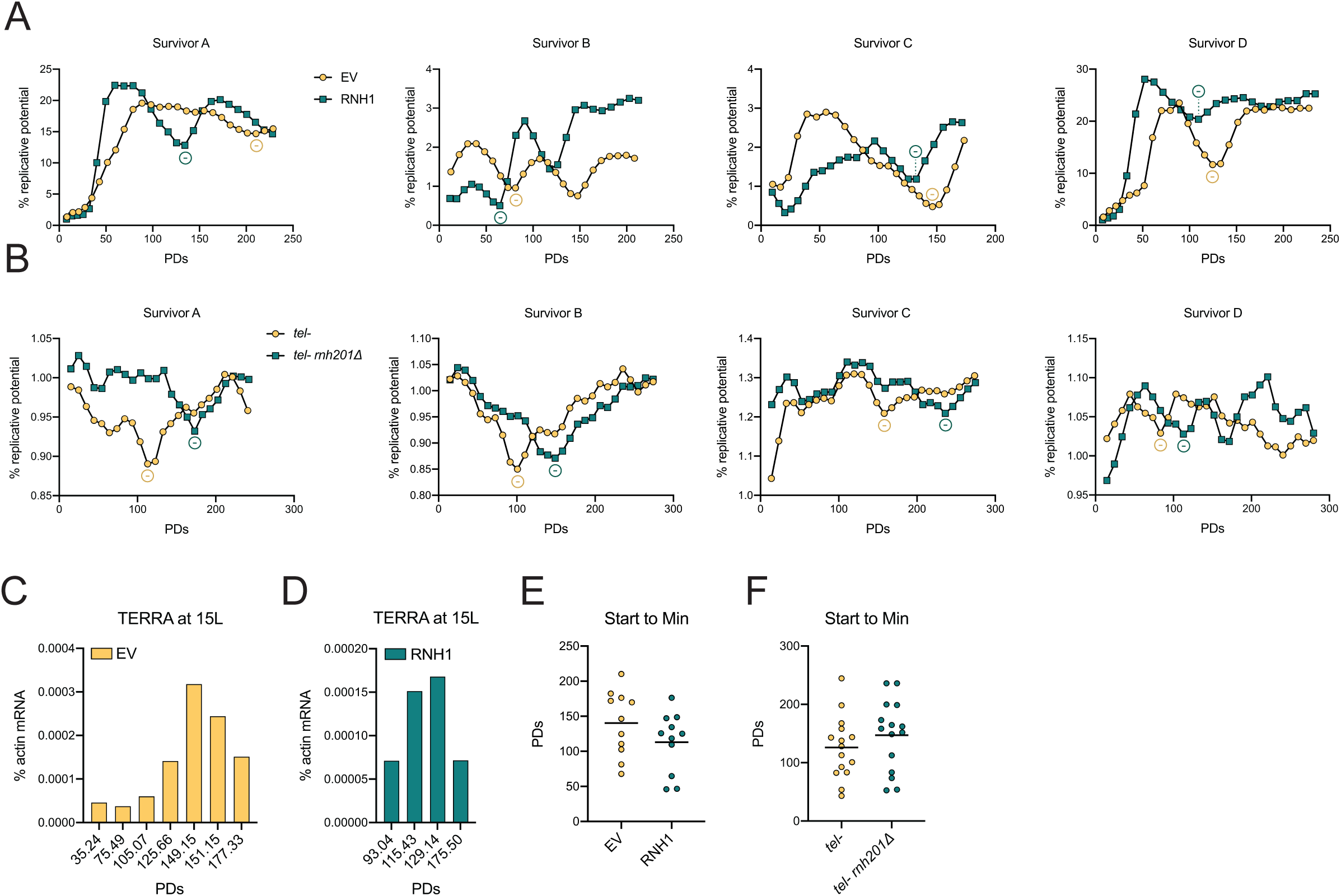
RNA-DNA hybrids regulate the rates of SAS. **(A)** Individual *tlc1Δ* survivor clones were transformed with an empty vector (EV) or an RNH1-overexpressing vector (RNH1) and propagated in liquid. The replicative potential was determined daily based on the optical density of each clonal survivor culture and plotted against the population doublings (PDs) values. Each “% replicative potential” value is divided by 100 for visualization reasons. “-” indicates the first local minimum (see Materials and Methods for more details). **(B)** The same approach in A) was used to monitor the changes of replicative potential in *est2Δ* or *tlc1Δ rnh201Δ* survivors transformed with an empty vector (*tel-rnh201Δ*) or a vector harboring the *RNH201* gene under its endogenous promoter (*tel-*). **(C)** The levels of TERRA from telomere 15L were monitored at the indicated population doublings (PDs) of survivor C from A) transformed with the EV. **(D)** TERRA levels were measured as in C) in the same survivor transformed with the RNH1-overexpressing vector. In both C) and D), the expression levels are represented as “% actin mRNA”. **(E)** The number of population doublings (PDs) was calculated between the starting point of the cell culture propagation and the first local minimum (Min) of the replicative potential in multiple survivors including those shown in A) to quantify the rate of SAS (n = 11) (see Materials and Methods for more details). **(F)** The same measurement as in E) was performed in survivors from B) along with multiple others (n = 15). In E and F, the mean is displayed, n=11/15.

The observation that type II survivors senesce was surprising considering that type II survivors are commonly referred to as having viability similar to that of wild-type cells^20–22,25^. The difference between the fluctuating replicative potential that we observe and the viability trend previously reported is that we propagated cells starting from single clones after survivors formation. We observed such clonal populations harbor telomeres with unique lengths (Figure 3), therefore the entire population is subjected to the same changes of replicative potential derived from variations of the telomere length. If a mixed population of cells present different initial lengths of the same telomere, the fraction of cells with the shortest length will undergo senescence but the effect on the general viability of the population will be masked by the cells with a longer length. We experienced this effect in the late stage of the population propagation, when the intensity of the fluctuations decreased probably due to the emergence of survivors with multiple telomere lengths after the shortening events.

In summary, telomere shortening leads to survivor associated senescence (SAS) and TERRA levels increase as survivors lose replicative potential.

### RNA-DNA hybrids influence the rates of SAS

We overexpressed RNase H1 to remove R-loops at telomeres in survivors, as previously described^28^. We observed accelerated rates of SAS, as demonstrated by the earlier occurrence of the lowest replicative potential values compared to the empty vector controls (Figure 4A, compare yellow to green). Accordingly, the amount of TERRA from telomere 15L with RNase H1 overexpression peaked at an earlier time point than in the EV control, presumably due to anticipated shortening of the chromosome end (Figure 4D, RNA extracted from survivor C panel (A)). To quantitatively analyze the effect of RNA-DNA hybrids deprivation on the rate of SAS, we compared the number of population doublings required to reach a local minimum, the lowest replicative potential value, within a neighborhood of 4 values from the start of the experiment between RNH1-overexpressing cells and the respective EV control (Figure 4E, Figure S4A). We counted PDs from the start of the liquid culture until the first minimum and all survivors were generated and transformed with identical timing in terms of PDs following survivor formation. We excluded minima in the first 2-3 days as growth it is frequently eratic in early population doublings following a transfer from solid to liquid media. Although the data was heterogeneous, in accordance with the nature of survivors, the number of PDs was reduced in survivors overexpressing RNase H1, indicating that, similar to replicative senescence, SAS appears to be accelerated in the absence of TERRA R-loops. Western blotting confirmed that Rnh1 was overexpressed throughout the entirety of the experiment when protein was extracted from clones at the end of the experiment (Supplementary Figure S4B) To confirm the role of RNA-DNA hybrids in defining the rate of SAS, we tested whether an increase of R-loops abundance had the opposite effect of RNase H1 overexpression as in the case during replicative senescence. We analyzed SAS rates in survivors depleted of *RNH201*, the catalytic subunit of RNase H2, and key enzyme responsible for removing RNA-DNA hybrids at telomeres^29^. We isolated telomerase negative type II survivors with an additional *RNH201* deletion and transformed them with a centromeric plasmid carrying *RNH201* under its endogenous promoter or an empty vector to obtain strains that were telomerase negative (*tel-*) or telomerase and *RNH201* negative (*tel-rnh201*Δ), respectively. When we propagated *tel-* and *tel-rnh201*Δ survivors, we observed that SAS was slightly delayed in cells deleted for *RNH201* compared to those with a functional enzyme (Figures 4B). Indeed, the number of population doublings required to reach the local minima of the replicative potential was increased in *rnh201*Δ survivors (Figure 4F). This suggests that accumulation of RNA-DNA hybrids counteracts senescence in survivors. Overall, these data indicate that type II survivors senesce in response to telomere shortening similar to pre-senescent cells, with TERRA RNA-DNA hybrids slowing the rate of SAS.

In pre-senescent cells, RNA-DNA hybrids counteract senescence by preventing telomere shortening^28^. Since this effect depends on *RAD52*, it has been assumed that RNA-DNA hybrids hinder telomere shortening by promoting telomeric HDR. Alternatively, and not mutually exclusive, RNA-DNA hybrids may impede telomere shortening by preventing end resection.

### RNA-DNA hybrids counteract telomere shortening in type II survivors

To understand if hybrids prevent telomere shortening in type II survivors, we monitored the dynamics of telomere 1L with a known starting length as it shortened and recombined in the presence and absence of TERRA RNA-DNA hybrids. We used survivor clones isolated during propagation experiments. In most of the cases, we used the survivor clone of Figure 3C and 3D, and started when the length of telomere 1L was approximately 350 bp. After transformation with an *RNH1*-overexpressing plasmid or an empty vector (EV), survivor cells were propagated for nearly 100 PDs and changes in the telomere length were examined by both Telomere-PCR (Figures 5A, B) and Southern blot (Figures 5C-E). As shown in Figure 5A and B, telomere 1L shortened at an average rate of 2.7 bp/PD in cells harboring an empty vector control. In the absence of RNA-DNA hybrids (*RNH1* overexpression), the shortening rate was 1.6 times higher, up to 6.7 bp/PD, and the minimal telomere length was almost 50% shorter (Figure 5B). To confirm these results, we also used Southern blotting, however, due to the large fragment sizes generated by Southern blotting we were not able to accurately calculate precise shortening rates, but rather used relative shortening rates (Figure 5C). Consistently, we observed an approximately 2-fold increase in shortening rate when RNA-DNA hybrids were removed through *RNH1* overexpression (Figures 5D, E). In agreement with faster rates of telomere loss, we observed an earlier elongation via HDR when *RNH1* was overexpressed (Figure 5C). These results suggest that, like in senescent cells, RNA-DNA hybrids counteract telomere shortening in type II survivors. Interestingly, as observed by Southern blotting, telomere 1L was able to undergo both type I and type II HDR regardless of the RNA-DNA hybrids abundance (Figure 5C), indicating either that extensive lengthening reactions in survivors do not require telomeric R-loops, or that overexpressed *RNH1* is not able to completely remove all hybrids. Overall, these data suggest a model where telomeric RNA-DNA hybrids modulate senescence in type II survivors by regulating the rate of telomere shortening.

**Figure 5.**
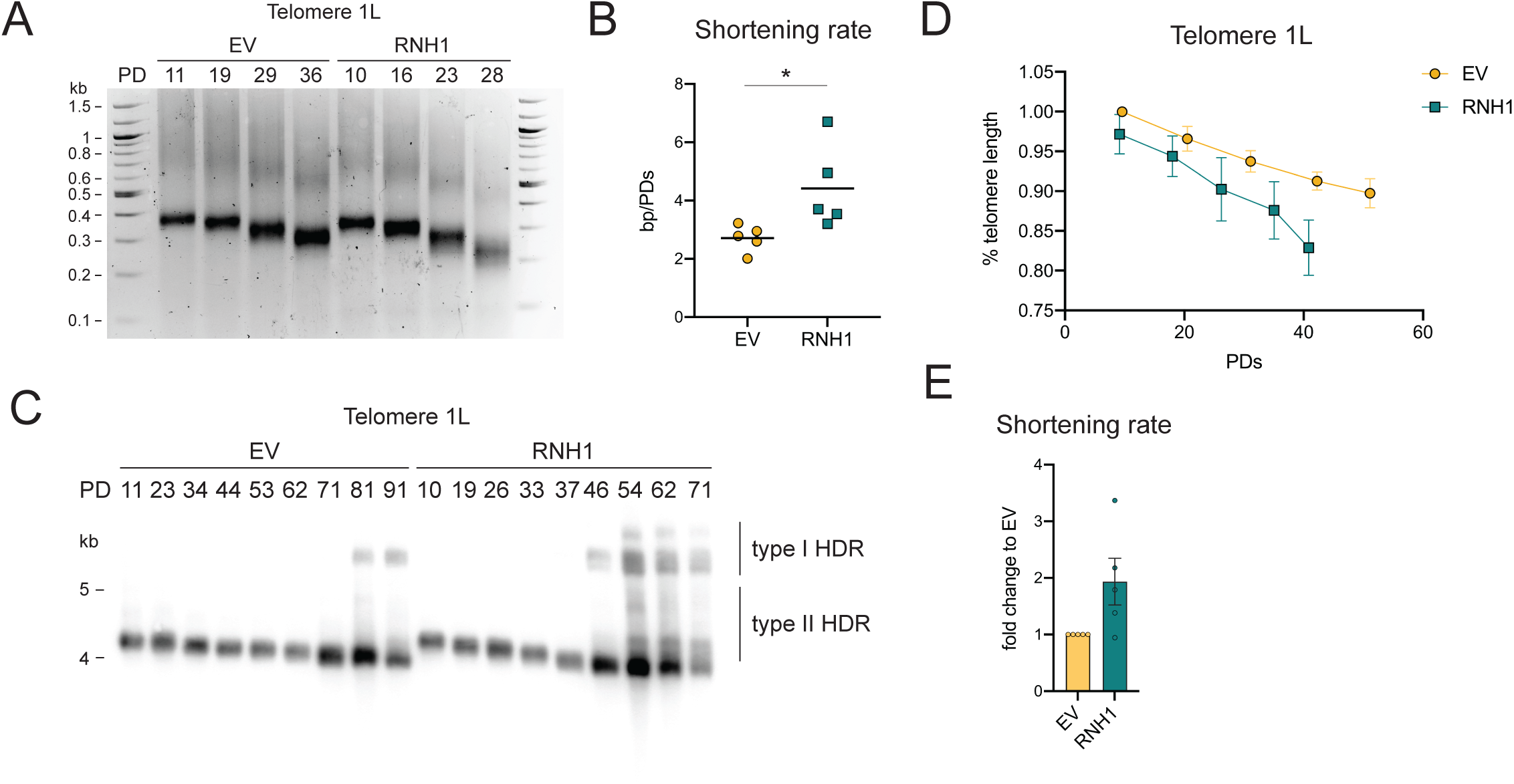
RNA-DNA hybrids counteract telomere shortening in type II survivors. **(A)** 1L telomere shortening was monitored by Telomere-PCR over multiple population doublings (PDs), obtained from the clonal propagation in Figure 3, transformed with an empty vector (EV) or an RNH1-overexpressing vector (RNH1). **(B)** The shortening rate of telomere 1L was derived by dividing the differences between two consecutive telomere lengths (in bp) by the differences between the corresponding population doublings (PD). The average of these “bp/PD” ratios was calculated for each of the five Telomere-PCR replicates and plotted. **(C)** Southern blot for telomere 1L as it shortens and undergoes HDR in one *tlc1Δ* survivor clone transformed with an empty vector (EV) or an RNH1-overexpressing plasmid (RNH1). The two types of HDR are shown. “PD” stays for population doublings. **(D)** The relative change of 1L telomere length was derived from the Southern blots of five survivor replicates and plotted as “% telomere length” of the first length in the EV strain (see Materials and Methods for more details). The “population doublings” (PDs) value plotted in the graph is the average among the five replicates. For visualization reasons, each value of “% telomere length” is divided by 100 in the graph. **(E)** The shortening rate of telomere 1L was derived from the linear regression of the shortening profile of each replicate represented in D) (see Materials and Methods for more details). To obtain the “fold change to EV” value, the linear regression of each RNH1 sample was divided by the one of the EV sample collected on the same day. The mean + SEM is displayed, n=5. In B, p values were calculated by an unpaired two-tailed Student’s *t*-test whereas in E, p values were calculated by an unpaired two-tailed Student’s *t*-test with Welch’s correction (*p<0.05).

In conclusion, our work identifies unexpected parallels between replicative senescence and the ALT phenotype (Figure 6). Upon telomerase loss, telomeres progressively shorten and cells gradually lose their proliferative potential. During this period, TERRA RNA-DNA hybrids seem to provide a protective role and prevent accelerated rates of senescence. When telomeres become critically short, TERRA levels strongly increase and this may trigger HDR and survivor formation. Surprisingly, we have now observed that once survivors are formed, the same trend continues in terms of telomere shortening and the TERRA RNA-DNA hybrid-mediated regulation of telomere shortening and senescence rates (Figure 6). These data, generated in yeast, may have important implications for human ALT cancers. Indeed, it will be interesting to see whether ALT cancer cells also senesce and whether they are susceptible to senolytic agents.

**Figure 6.**
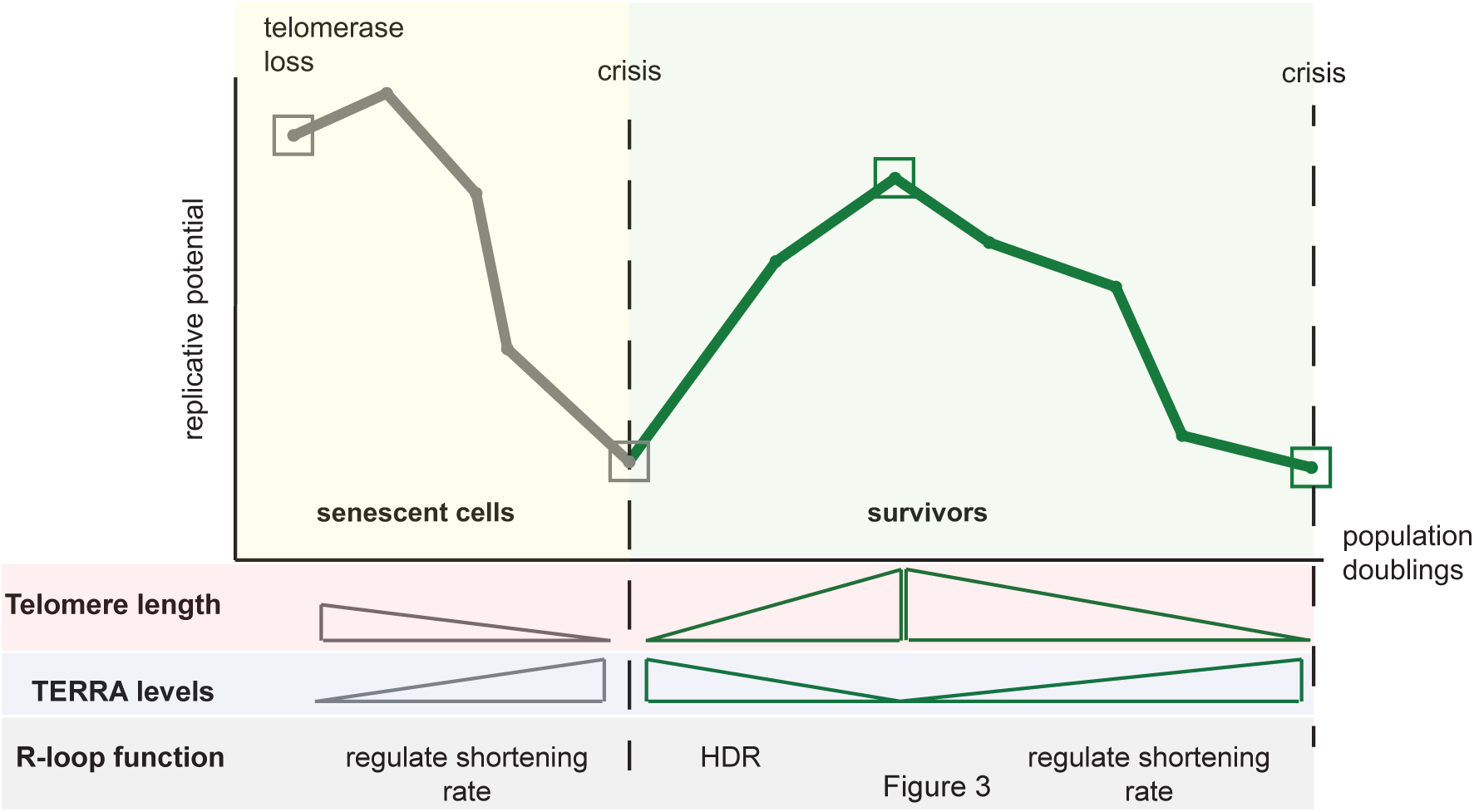
TERRA as a regulator of senescence and SAS. Upon telomerase loss, telomeres shorten. The continued presence of TERRA RNA-DNA hybrids at telomeres ensures that senescence is not accelerated. This may be accomplished through HDR-mediated extension or by preventing resection. As telomeres get critically short, TERRA and RNA-DNA hybrids get stabilized. This increases the likelihood of HDR events and the formation of survivors (yellow shading – replicative senescence). Upon survivor formation (green shading), telomeres are extensively elongated and TERRA levels are decreased. However, survivors also experience telomere shortening and undergo replicative senescence, we refer to this as survivor associated senescence (SAS). During this process, TERRA RNA-DNA hybrids prevent telomere shortening and regulate SAS rates. Eventually, at critically short telomeres TERRA and hybrids become stabilized which again increases the chance of HDR-mediated elongation.

## Data Availability

No NGS or proteomics data sets were generated during this study.

## Acknowledgements

We thank the members of the Luke group for helpful discussions, Diego Bonetti for generating part of the survivor strains and Kathy Friedman for the *TLC1* plasmid. Support by the IMB Media Lab, Flow cytometry, and Protein production core facilities is gratefully acknowledged.

## References

1. De Lange, T. How telomeres solve the end-protection problem. Science 326, 948–952 (2009).

2. Wellinger, R. J. & Zakian, V. A. Everything you ever wanted to know about Saccharomyces cerevisiae telomeres: Beginning to end. Genetics 191, 1073–1105 (2012).

3. Teixeira, M. T. Saccharomyces cerevisiae as a model to study replicative senescence triggered by telomere shortening. Front. Oncol. 3, 1–16 (2013).

4. Claussin, C. & Chang, M. The many facets of homologous recombination at telomeres. Microb. Cell 2, 308–321 (2015).

5. Lingner, J., Cooper, J. P. & Cech, T. R. Telomerase and DNA end replication: No longer a lagging strand problem? Science 269, 1533–1534 (1995).

6. Lundblad, V. & Szostak, J. W. A mutant with a defect in telomere elongation leads to senescence in yeast. Cell 57, 633–643 (1989).

7. Soudet, J., Jolivet, P. & Teixeira, M. T. Elucidation of the DNA end-replication problem in saccharomyces cerevisiae. Mol. Cell 53, 954–964 (2014).

8. Lendvay, T. S., Morris, D. K., Sah, J., Balasubramanian, B. & Lundblad, V. Senescence Mutants of Saccharomyces Cerevisiae With a Defect in Telomere Replication Identify Three Additional EST Genes. Genetics 144, 1399–1412 (1996).

9. Collado, M., Blasco, M. A. & Serrano, M. Cellular Senescence in Cancer and Aging. Cell 130, 223–233 (2007).

10. Campisi, J. Aging, cellular senescence, and cancer. Annu. Rev. Physiol. 75, 685–705 (2013).

11. Greider, C. W. & Blackburn, E. H. Identification of a specific telomere terminal transferase activity in tetrahymena extracts. Cell 43, 405–413 (1985).

12. Wright, W. E., Piatyszek, M. A., Rainey, W. E., Byrd, W. & Shay, J. W. Telomerase activity in human germline and embryonic tissues and cells. Dev. Genet. 18, 173–179 (1996).

13. Wright, D., Gibbons, W. & Lanzendorf, S.. Characterization of Telomerase Activity in the Human Oocyte and Preimplantation Embryo. Fertil. Steril. 74, S67 (2000).

14. Kim, N. W. et al. Specific association of human telomerase activity with immortal cells and cancer. Science 266, 2011–2015 (1994).

15. Shay, J. W. Role of telomeres and telomerase in aging and cancer. Cancer Discov. 6, 584–593 (2016).

16. Shay, J. W. & Bacchetti, S. A survey of telomerase activity in human cancer. Eur. J. Cancer Part A 33, 787–791 (1997).

17. Bryan, T. M., Englezou, A., Dalla-Pozza, L., Dunham, M. A. & Reddel, R. R. Evidence for an alternative mechanism for maintaining telomere length in human tumors and tumor-derived cell lines. Nat. Med. 3, 1271–1274 (1997).

18. Shay, J. W., Reddel, R. R. & Wright, W. E. Cancer and Telomeres— An ALTernative to Telomerase. Science 336, 1388–1390 (2012).

19. Singer, M. S. & Gottschling, D. E. TLC1: Template RNA component of Saccharomyces cerevisiae telomerase. Science 266, 404–409 (1994).

20. Lundblad, V. & Blackburn, E. H. An alternative pathway for yeast telomere maintenance rescues est1-senescence. Cell 73, 347–360 (1993).

21. Teng, S. C. & Zakian, V. A. Telomere-telomere recombination is an efficient bypass pathway for telomere maintenance in Saccharomyces cerevisiae. Mol. Cell. Biol. 19, 8083–93 (1999).

22. Teng, S. C., Chang, J., McCowan, B. & Zakian, V. a. Telomerase-independent lengthening of yeast telomeres occurs by an abrupt Rad50p-dependent, Rif-inhibited recombinational process. Mol. Cell 6, 947–52 (2000).

23. Lydeard, J. R., Jain, S., Yamaguchi, M. & Haber, J. E. Break-induced replication and telomerase-independent telomere maintenance require Pol32. Nature 448, 820–823 (2007).

24. Kockler, Z. W., Comeron, J. M. & Malkova, A. A unified alternative telomere-lengthening pathway in yeast survivor cells. Mol. Cell 81, 1–14 (2021).

25. Grandin, N. & Charbonneau, M. Telomerase- and Rad52-Independent Immortalization of Budding Yeast by an Inherited-Long-Telomere Pathway of Telomeric Repeat Amplification. Mol. Cell. Biol. 29, 965–985 (2009).

26. Azzalin, C. M., Reichenbach, P., Khoriauli, L., Giulotto, E. & Lingner, J. Telomeric Repeat Containing RNA and RNA Surveillance Factors at Mammalian Chromosome Ends. Science 318, 798–801 (2007).

27. Luke, B. et al. The Rat1p 5′ to 3′ Exonuclease Degrades Telomeric Repeat-Containing RNA and Promotes Telomere Elongation in Saccharomyces cerevisiae. Mol. Cell 32, 465–477 (2008).

28. Balk, B. et al. Telomeric RNA-DNA hybrids affect telomere-length dynamics and senescence. Nat. Struct. Mol. Biol. 20, 1199–1206 (2013).

29. Graf, M. et al. Telomere Length Determines TERRA and R-Loop Regulation through the Cell Cycle. Cell 170, 72-85.e14 (2017).

30. Bettin, N., Oss Pegorar, C. & Cusanelli, E. The Emerging Roles of TERRA in Telomere Maintenance and Genome Stability. Cells 8, 246 (2019).

31. Arora, R. et al. RNaseH1 regulates TERRA-telomeric DNA hybrids and telomere maintenance in ALT tumour cells. Nat. Commun. 5, 5220 (2014).

32. Arora, R. & Azzalin, C. M. Telomere elongation chooses TERRA ALTernatives. RNA Biol. 12, 938–941 (2015).

33. Silva, B. et al. FANCM limits ALT activity by restricting telomeric replication stress induced by deregulated BLM and R-loops. Nat. Commun. 10, 1–16 (2019).

34. Episkopou, H. et al. Alternative Lengthening of Telomeres is characterized by reduced compaction of telomeric chromatin. Nucleic Acids Res. 42, 4391–4405 (2014).

35. Misino, S., Bonetti, D., Luke-Glaser, S. & Luke, B. Increased TERRA levels and RNase H sensitivity are conserved hallmarks of post-senescent survivors in budding yeast. Differentiation 100, (2018).

36. Lovejoy, C. A. et al. Loss of ATRX, genome instability, and an altered DNA damage response are hallmarks of the alternative lengthening of Telomeres pathway. PLoS Genet. 8, 12–15 (2012).

37. Azzalin, C. M., Reichenbach, P., Khoriauli, L., Giulotto, E. & Lingner, J. Telomeric repeat-containing RNA and RNA surveillance factors at mammalian chromosome ends. Science 318, 798–801 (2007).

38. Schoeftner, S. & Blasco, M. A. Developmentally regulated transcription of mammalian telomeres by DNA-dependent RNA polymerase II. Nat. Cell Biol. 10, 228–236 (2008).

39. Silva, B., Arora, R., Bione, S. & Azzalin, C. M. TERRA transcription destabilizes telomere integrity to initiate break-induced replication in human ALT cells. Nat. Commun. 12, 1–12 (2021).

40. Hu, Y. et al. RNA-DNA hybrids support recombination-based telomere maintenance in fission yeast. Genetics 213, 431–437 (2019).

41. Yu, T.-Y., Kao, Y. & Lin, J.-J. Telomeric transcripts stimulate telomere recombination to suppress senescence in cells lacking telomerase. Proc. Natl. Acad. Sci. 111, 3377–3382 (2014).

42. Porro, A., Feuerhahn, S., Reichenbach, P. & Lingner, J. Molecular Dissection of Telomeric Repeat-Containing RNA Biogenesis Unveils the Presence of Distinct and Multiple Regulatory Pathways. Mol. Cell. Biol. 30, 4808–4817 (2010).

43. Flynn, R. L. et al. Alternative lengthening of telomeres renders cancer cells hypersensitive to ATR inhibitors. Science 347, 273–277 (2015).

44. Vohhodina, J. et al. BRCA1 binds TERRA RNA and suppresses R-Loop-based telomeric DNA damage. Nat. Commun. 12, 3542 (2021).

45. Teixeira, M. T., Arneric, M., Sperisen, P. & Lingner, J. Telomere length homeostasis is achieved via a switch between telomerase-extendible and -nonextendible states. Cell 117, 323–335 (2004).

46. Fallet, E. et al. Length-dependent processing of telomeres in the absence of telomerase. Nucleic Acids Res. 42, 3648–3665 (2014).

47. Hardy, J., Churikov, D., Géli, V. & Simon, M.-N. Sgs1 and Sae2 promote telomere replication by limiting accumulation of ssDNA. Nat. Commun. 2014 51 5, 1–13 (2014).

48. Craven, R. J. & Petes, T. D. Dependence of the regulation of telomere length on the type of subtelomeric repeat in the yeast Saccharomyces cerevisiae. Genetics 152, 1531–1541 (1999).

49. Marcand, S., Gilson, E. & Shore, D. A protein-counting mechanism for telomere length regulation in yeast. Science 275, 986–990 (1997).

50. Marcand, S., Brevet, V. & Gilson, E. Progressive cis-inhibition of telomerase upon telomere elongation. EMBO J. 18, 3509–3519 (1999).

