## Supplementary Data for "TERRA increases at short telomeres in yeast survivors and regulates survivor associated senescence (SAS)"

**Supplementary Figures, Misino et al**

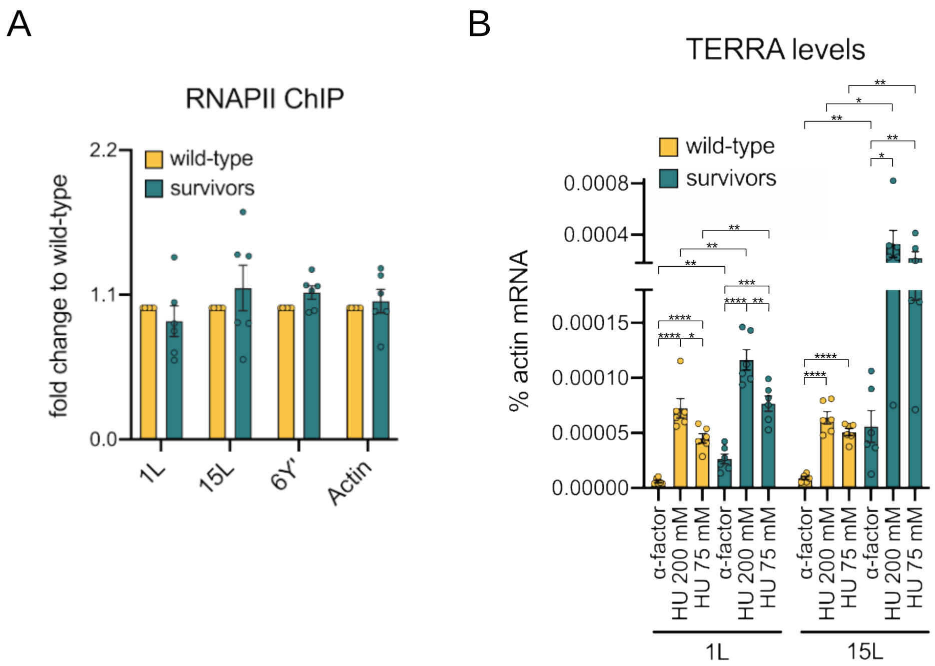


**Figure S1. TERRA cell cycle regulation is upheld**

**(A)** ChIP of RNA pol II at 1L, 15L and 6Y' telomeres in wild-type and survivor cells. The "% input" values of all samples were normalized by the ones of wild-type cells to obtain the final "fold change to wild-type". The actin locus was used as a positive control.

**(B)** TERRA levels were measured at telomere 1L and 15L in *tlc1Δ* survivors and wild-type cells arrested in G1 (a-factor), early S-phase (HU 200 mM) and late S-phase (HU 75 mM). The amount of TERRA is expressed as "% actin mRNA". HU indicates hydroxyurea.

Mean + SEM is displayed, n=6. p values were calculated by an unpaired two-tailed Student's *t*-test (*p<0.05, **p<0.01, ***p<0.001, ****p<0.0001).

A

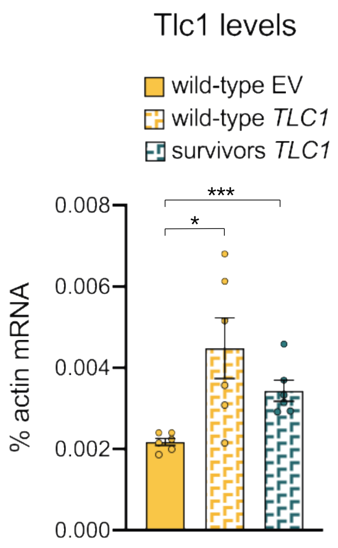


**Figure S2. *TLC1* expression in wild type and survivor cells**

**(A)** *TLC1* expression levels were monitored in wild-type cells transformed with an empty vector (EV) and in wild-type cells and survivors transformed with a vector expressing the gene under its endogenous promoter (*TLC1*). The expression levels are represented as "% actin mRNA". Mean + SEM is displayed, n=6. p values were calculated by an unpaired two-tailed Student's *t*-test (*p<0.05, ***p<0.001).

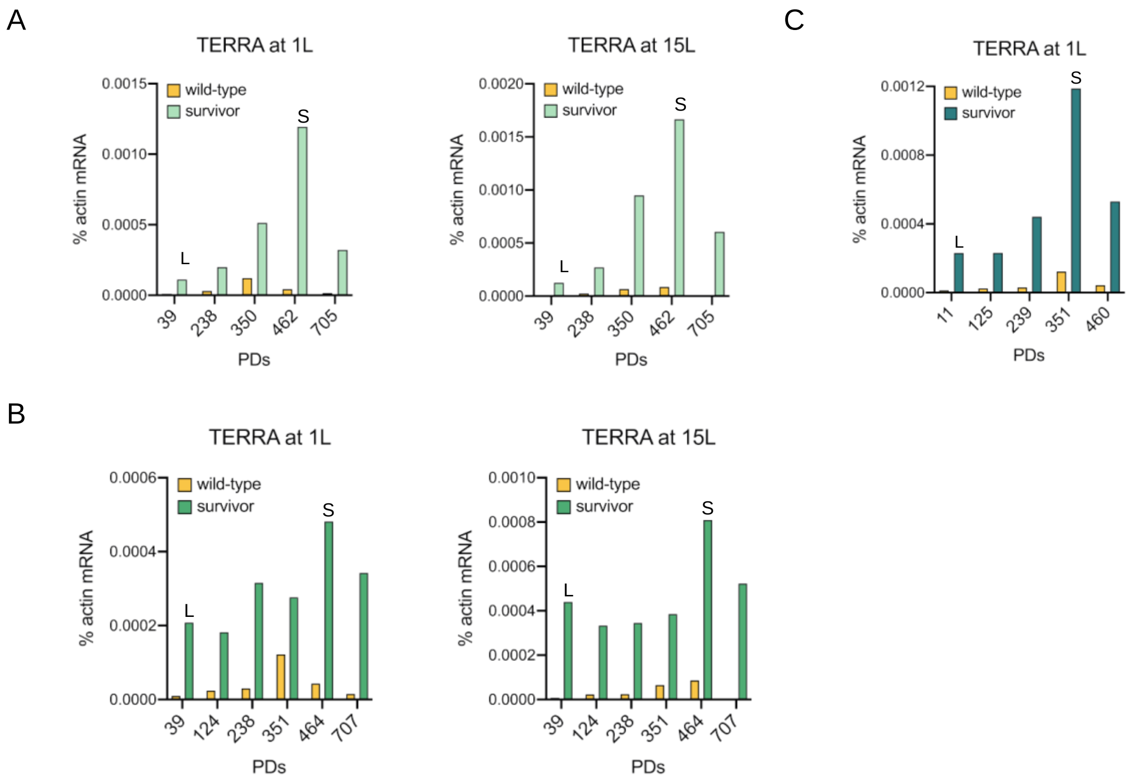


**Figure S3. TERRA levels increase at short telomeres in survivors**

**(A, B, C)** TERRA levels were monitored in 3 independent survivor clones. “L” and “S” indicate when telomeres were long and when they were shortened as followed by Southern blots. The levels of TERRA deriving from these telomeres are plotted. 15L TERRA levels are not reported for clone C as the telomere underwent an early recombination event and acquired a Y´element, hence complicating interpretation of the result.

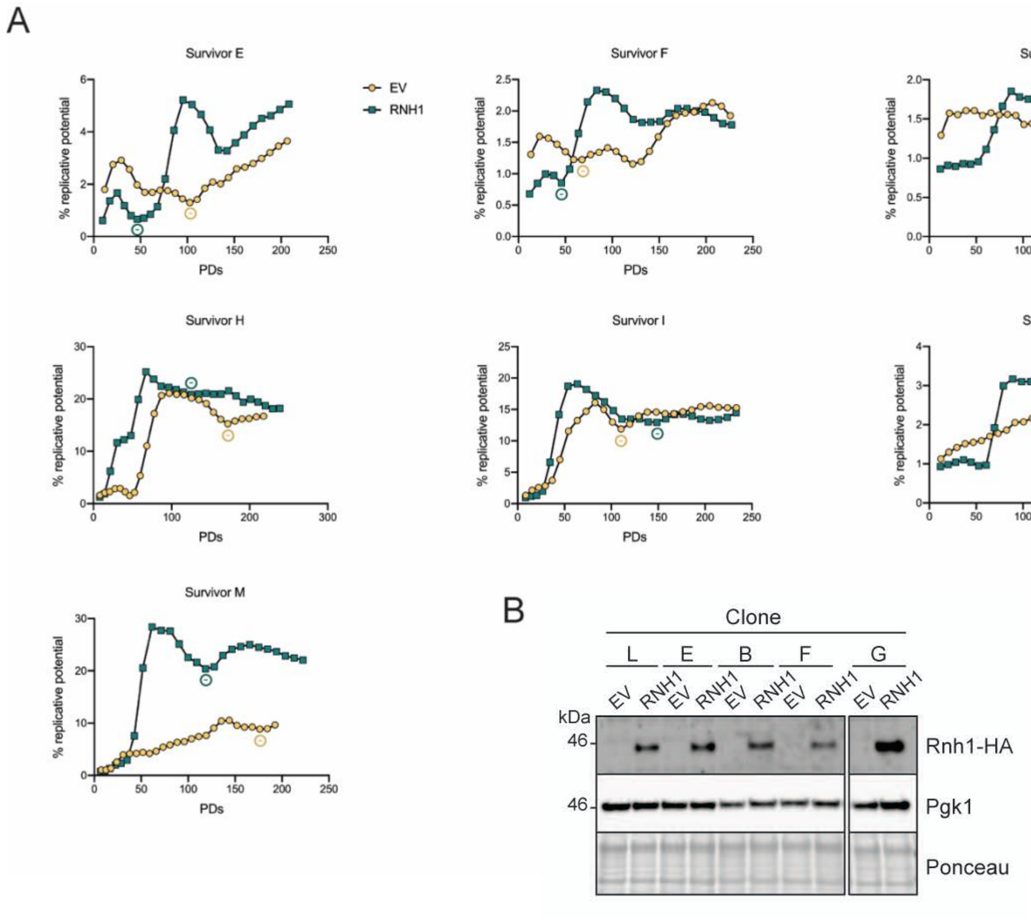


**Figure S4. SAS rates are increased with RNase H1 overexpression**

**(A)** All growth curves that contributing to the data in Figure 4E and performed in a manner similar to Figures 4A and 4B.

(**B)** Western blot showing the overexpression of Rnh1-HA in survivors Figure 4A. Pgk1 and Ponceau staining were used as loading control. These extracts were performed on the last day of the passaging experiment.
